# Etrinabdione (VCE-004.8), a B55α activator, promotes angiogenesis and arteriogenesis in critical limb ischemia

**DOI:** 10.1101/2024.04.26.591240

**Authors:** Adela García-Martín, María E. Prados, Isabel Lastres-Cubillo, Francisco J. Ponce-Diaz, Laura Cerero, Martin Garrido-Rodríguez, Carmen Navarrete, Rafael Pineda, Ana B. Rodríguez, Ignacio Muñoz, Javier Moya, Antonella Medeot, José A. Moreno, Antonio Chacón, José García-Revillo, Eduardo Muñoz

## Abstract

**Background:** Vasculogenic therapies explored for the treatment of peripheral artery disease (PAD) have encountered minimal success in clinical trials. Addressing this, B55α, an isoform of protein phosphatase 2A (PP2A), emerges as pivotal in vessel remodeling through activation of hypoxia-inducible factor 1α (HIF-1α). This study delves into the pharmacological profile of VCE-004.8 (Etrinabdione) and evaluates its efficacy in a preclinical model of critical limb ischemia, with a focus on its potential as a PP2A/B55α activator to induce angiogenesis and arteriogenesis.

**Methods:** Vascular endothelial cells were used for *in vitro* experiments. Aorta ring assay was performed to explore sprouting activity. Matrigel plug-in assay was used to assess the angiogenic potential. Critical limb ischemia (CLI) in mice was induced by double ligation in the femoral arteria. Endothelial vascular and fibrotic biomarkers were studied by immunohistochemistry and qPCR. Arteriogenesis was investigated by microvascular casting and micro-CT. Proteomic analysis in vascular tissues was analyzed by LC-MS/MS. *Ex-vivo* expression of B55α and biomarkers were investigated in artery samples from PAD patients.

**Results:** VCE-004.8 exhibited the ability to induce B55α expression and activate the intersecting pathways B55α/AMPK/Sirtuin 1/eNOS and B55α/PHD2/HIF-1α. VCE-004.8 prevented OxLDL and H_2_O_2_-induced cytotoxicity, senescence, and inflammation in endothelial cells. Oral VCE-004.8 increased aorta sprouting *in vitro* and angiogenesis *in vivo*. In CLI mice VCE-004.8 improved collateral vessel formation and induced endothelial cells proliferation, angiogenic gene expression and prevented fibrosis. The expression of B55α, Caveolin 1 and Sirtuin-1 is reduced in arteries from CLI mice and PAD patient, and the expression of these markers was restored in mice treated with VCE-004.8.

**Conclusions:** The findings presented in this study indicate that Etrinabdione holds promise in mitigating endothelial cell damage and senescence, while concurrently fostering arteriogenesis and angiogenesis. These observations position Etrinabdione as a compelling candidate for the treatment of PAD, and potentially other cardiovascular disorders.

**Novelty and Significance:** *What Is Known?:*

- The phosphatase PPA2/B55α stabilizes endothelial cells (ECs) in response to cell stress conditions, thereby protecting ECs from apoptosis and promoting angiogenesis.
- Etrinabdione (VCE-004.8) functions as a potent activator of PPA2/B55α inducing PHD2 dephosphorylation at ser125 and fostering HIF activation.
- VCE-004.8 prevents vascular damage in preclinical models of systemic and cardiac fibrosis and alleviates blood-brain barrier disruption in neuroinflammatory conditions.
- VCE-004.8 is also a dual agonist of PPARγ and CB_2_ receptors and shows antiinflammatory activity.
- Oral VCE-004.8 has meet the primary endpoints of safety and tolerability in a Phase IIa clinical trial with systemic sclerosis patients (clinicaltrial.gov: NCT03745001).

*What New Information Does This Article Contribute?:*

- Etrinabdione induces HIF-1α expression in endothelial cells through a novel pathway that potentially involves two axes: B55α/PHD2 and B55α/AMPK/Sirt1 signaling that may converge on HIF stabilization.
- Etrinabdione prevented endothelial cell damage and senescence, while inducing arteriogenesis and angiogenesis in CLI mice.
- In arteries of patients with PAD and in CLI mouse models, the expression levels of B55α, Caveolin 1, and Sirtuin 1 are diminished. However, treatment with Etrinabdione specifically in CLI mice prompts an increase in the levels of these proteins.
- Etrinabdione triggers neovascularization and angiogenesis specifically within hypoxic tissue in a critical ischemia model, with no impact on healthy tissue.

## INTRODUCTION

Peripheral arterial disease (PAD) is one of the major leading cause of atherosclerotic cardiovascular morbidity^1^. Experimental approaches to the treatment of PAD have included the use of angiogenic factors as a recombinant protein or as gene therapy as well as stem cell therapy^2^ but, neither of these strategies achieved significant benefits in clinical trials. Hence, there is an urgent demand for novel pharmacological targets and therapies capable of preventing endothelial damage and promoting angiogenesis and arteriogenesis to enhance blood flow in extremities.

The protein phosphatase 2A (PP2A) belong to a family of serine-threonine phosphatases, which are formed by a catalytic C subunit, a scaffold A subunit, and a regulatory B subunit, which confers specificity of PP2A for selective proteins substrates^3^. B55α (PPP2R2A) is one of the regulatory subunits that has been shown to directly dephosphorylates prolyl hydroxylase 2 (PHD2) on Ser125, resulting in a further reduction of its activity that ultimately stabilized and accumulated the hypoxia inducing factor 1α (HIF-1α)^4^. HIF-1, a heterodimeric basic-helix-loop-helix-paste protein that responds to oxygen tension, is activated by severe and mild hypoxia. Under hypoxic conditions, HIF-1α accumulates in the nucleus and dimerizes with HIF-1β to activate a plethora of genes including angiogenic factors such as vascular endothelial growth factor (VEGF), fibroblast growth factor (FGF) and angiopoietin, as well as an important number of genes involved in vascular and tissue remodeling.^5,6^ HIF-1α is also activated by therapeutic preconditioning hypoxia and by hypoxia mimetic small compounds.^7,8^ More recently, it has been shown that B55α stabilizes endothelial cells (ECs) in response to cell stress conditions, thereby protecting ECs from apoptosis and promoting angiogenesis. Thus, PP2A/B55α activators represent a new class of hypoxia mimetic compounds with potential in vessel remodeling, angiogenesis and endothelial cell protection^9^.

The histone deacetylase Sirtuin 1 (Sirt1) belong to the family of Sirtuins, which are nicotinamide adenine dinucleotide (NAD^+^)-dependent enzymes with multiple metabolic functions. The expression of Sirt1 is reduced with aging both in animal models and in humans, and a reduced expression is thought to be involved in age-related cardiovascular diseases. Indeed, Sirt1, which is activated by AMPK, is a critical modulator of the vascular function^10^ and endothelial senescence is regarded as a prominent precursor of cardiovascular diseases including PAD and critical limb ischemia (CLI).^11^ The key role of Sirt1 in the regulation of vascular homeostasis is mediated through complex cellular mechanisms involved in the promotion of vasodilatory and regenerative functions at ECs, endothelial progenitor cells, and smooth muscle cells (SMC) levels.^12^

We have recently identified the cannabidiol aminoquinone VCE-004.8 (also known as Etrinabdione and EHP-101), as a PP2A/B55α activator that induces HIF-1α and HIF-2α stabilization.^13^ In addition, VCE-004.8 is a dual activator of peroxisome proliferator-activated receptor-γ (PPARγ) and cannabinoid type 2 receptors (CB_2_R).^14^ VCE-004.8 showed efficacy in preclinical models of traumatic brain injury,^15^ multiple sclerosis^13^, bleomycin- and angiotensin-induced models of dermal, lung, cardiac, and kidney fibrosis^14, 16^, metabolic syndrome^17^ and stroke^18^.

Etrinabdione (VCE-004.8) has finalized the stage 1 of a clinical Phase IIa trial in Systemic Sclerosis patients (Clinical Trials.gov: NCT04166552), and it had acceptable safety profile and was well-tolerated. Herein we have further investigated additional pharmacological attributes of VCE-004.8 and explored its efficacy in a model of CLI.

## METHODS

Please see the Supplemental Material for detailed methods on cell cultures, transient transfections and luciferase assays, western blot analysis, measurement of NO (nitric oxide), Sirtuin 1 activity, NAD^+^ /NADH ratio, senescence assay, aortic ring assay, *in vivo* angiogenesis, in *vivo* perfusion fixation and tissue processing procedures, confocal immunofluorescence microscopy, qPCR, and proteomic analysis). All experimental protocols followed the guidelines of animal care set by the EU guidelines 86/609/EEC, the Ethic Committee on Animal Experimentation at the University of Córdoba (Spain) and the Andalusian Regional Committee for Animal Experimentation approved all the procedures described in this study (31/03/2022/057). Human biopsies from patients were obtained by the Cardiovascular Surgery Unit of Hospital Universitario Reina Sofia of Cordoba, Spain (RSUH) and the protocol approved from the Ethics Committee of RSUH (reference: HIP-CI:1.0-01/06/2023).

### Data Availability

The data that support the findings of this study are available from the corresponding author upon request.

### Test compound

VCE-004.8 (also called Etrinabdione and EHP-101) [(1′R,6′R)−3-(Benzylamine)−6-hydroxy-3′-methyl-4-pentyl-6′-(prop-1-en-2-yl) [1,1′bi(cyclohexane)]− 2′,3,6-triene-2,5-dione)] was produced under GMP conditions and provided by VivaCell Biotechnology España, Spain). For animal use VCE-004.8 was dissolved in sesame oil/Maisine CC^®^ (50%/50%) and applied by oral gavage.

### Critical limb ischemia (CLI) mouse model

Male C57BL/6 mice aged 10-12 weeks were divided into 2 groups (CLI group and sham group). In the CLI group the femoral artery and its side branches were double ligated with 6–0 silk sutures immediately distal to the inguinal ligament and proximal to the popliteal bifurcation. Femoral nerves were carefully preserved. The wound was irrigated with sterile saline and then the overlying skin was closed using 4–0 vicryl sutures. Post-operative pain was reduced using Lignocaine. A similar surgery without ligation of the femoral artery was performed on the sham controls. The CLI group was administered VCE-004.8 (20 mg/kg) by oral gavage every day until the end of the study and the sham control group was treated with vehicle (Corn oil/Maisine^®^ CC (50/50 (v/v)). At 10 and 28 days after ligation, the whole limb and muscles were harvested for study.

### Vascular casting

Mice were anesthetized and injected 1000 UI of heparin (i.p.). After 10 min, the mice were euthanized, the thorax was opened, and the aorta exposed. A catheter was inserted into the descending aorta and fixed with sutures. Then mice were perfused with PBS (37 °C, 80 ml) containing heparin, nitroglycerin, adenosine and sodium nitroprusside to remove the blood and enhanced the vasodilatation of the vessels. Microfil (MV-112 [white]; Flow Tech Inc, South Windsor, CT, USA) was perfused and then filled the entire vascular bed. The Microfil polymerized overnight at 4°C, and the samples were harvested and clarified in graded glycerol solutions (40%–100% glycerol in water; glycerol increased by 20% at 24-hour intervals). The clarified specimens were viewed on a dissecting microscope and Microfil retained in the vasculature was investigated by μ-CT (X-ray tomography).

### Image acquisition μ-CT

After Microfilm polymerization, the animals were hold inside a Bruker SkyScan 1172 high resolution microtomography machine (Bruker micro-CT, Kontich, Belgium). The X-ray source was set to a voltage of 50 kV and a current of 498 μA with a 0.5-mm Al filter in the beam path with an angular increment of 0.3°. Data were transferred to a computer with NRecon, CTAn, CT Vol software, (Bruker), ImageJ (http://rsb.info.nih.gov/ij/) and 3DSlicer v.3.4.0 open-source software (https://pubmed.ncbi.nlm.nih.gov/22770690/) was using to analyze vessel numbers, diameter, area, and volume, as well as arterial density.

### Laser capture microdissection (LCM) targeted mass spectroscopy

LCM and subsequent molecular analysis were carried out on slides immuno-stained using anti-α-Smooth Muscle Actin (α-SMA) antibody of gastrocnemius muscles frozen sections. The vessel marked with immunofluorescence α-SMA were dissected with ZEISS PALM Microbeam micro dissected and collected for mass spectroscopy analysis.

### Proteomics data analysis

To generate Figure 7B, we employed the log transformed intensities for proteins annotated with the Gene Ontology term ‘Angiogenesis’ (GO: 0001525). The mass spectrometry proteomics data have been deposited to the ProteomeXchange Consortium via the PRIDE partner repository with the dataset identifier PXD050360.

### Statistical Analysis

Data were expressed as the mean ± SEM or SD. One-way analysis of variance (ANOVA) followed by the Tukeýs post-hoc test for parametric analysis or Kruskal-Wallis post-hoc test in the case of non-parametric analysis tests will be used to determine the statistical significance. The level of significance will be set at p˂ 0.05. Statistical analyses will be performed using GraphPad Prism version 9 (GraphPad, San Diego, CA, USA).

## RESULTS

### VCE-004.8 induces B55α/PP2A in endothelial vascular cells

We have demonstrated that VCE-004.8, a hypoxia-mimetic cannabidiol aminoquinone derivative, dephosphorylates PHD2 at ser-125 via a PP2A/B55α-dependent pathway, leading to the induction of HIF-1α expression^13^. In this study, we further reveal that B55α expression in EA-hy926 cells is primarily localized perinuclearly and is significantly upregulated following exposure to VCE-004.8 (Fig. 1A). As anticipated, silencing B55α expression with siRNA in EA-hy926 cells abrogated the VCE-004.8-induced stabilization of HIF-1α (Fig. 1B). To investigate the functional impact of VCE-004.8 and HIF-1α on cell viability, we subjected the cells to H_2_O_2_ treatment in the presence or absence of the compound. VCE-004.8 greatly prevented H_2_O_2_-cytotoxicity in a concentration-dependent manner (Fig. 1C), which was significantly counteracted by YC-1, a HIF-1α inhibitor (Fig. 1D). Moreover, the cytoprotective effect of VCE-004.8 was evident in cells exposed to both high glucose and OxLDL (Fig. 1S). Subsequently, to explore the anti-inflammatory properties of VCE-004.8, EA-hy926 cells were incubated with proinflammatory cytokines, and the expression of various markers was analyzed via confocal microscopy. VCE-004.8 attenuated IL-1β+TNFα-induced VCAM-1 expression and restored the expression of tight-junction proteins ZO-1 and CLD1 in IL-6/TNFα-treated cells. Notably, tight-junction proteins play a crucial role in maintaining the integrity of vascular endothelial cells^19^. Furthermore, VCE-004.8 also induced the expression of B55α under inflammatory conditions (Fig. 2S).

**Figure 1.**
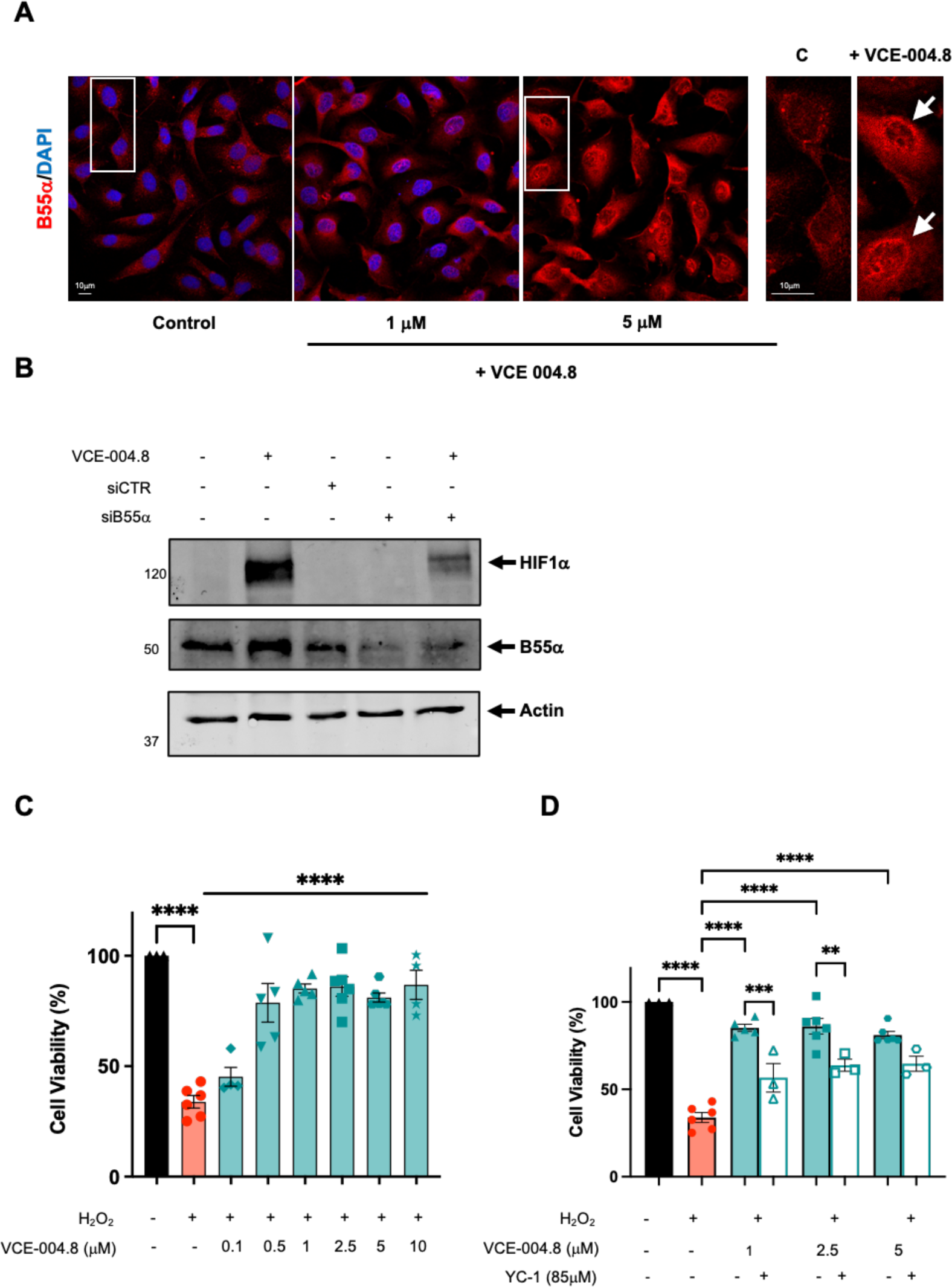
VCE-004.8 induces B55α/PP2A expression and activates HIF-1α in endothelial EA.hy926 cells. **A,** Representative images of B55α expression in EA.hy926 cells untreated and treated with VCE-004.8 for 24 h. Magnified region (marked by the white rectangle) shows preferential perinuclear expression of B55α as indicated by white arrows. **B,** EA.hy926 cells were transfected with siB55α or scrambled siRNAs (siCTR) and 48 h later treated with VCE-004.8 for 3 h. B55α and HIF-1α protein expression was analyzed by immunoblotting. **C,** Effect of VCE-004.8 on cell viability. EA.hy926 cells were pre-incubated with increasing concentrations of VCE-004.8 during 1 h and then were exposed to H_2_O_2_ for 24 h. Cell viability was calculated by MTT assay. The data passed the normality test performed by the Kolmogorov-Smirnov test. P value is calculated using 1-way ANOVA followed by Tukey post hoc multiple comparisons to compare between the groups (n>4 independent experiments per group), the results are shown as mean ± SEM. **D**, The cytoprotection mediated by VCE-004.8 is partially dependent on HIF-1α activation. EA.hy926 cells were pretreated with VCE-004.8 in the absence and the presence of the HIF-1α inhibitor YC-1 and then exposed to H_2_O_2_. Cell viability was calculated by MTT assay. The data passed the normality test performed by the Shaphiro-Wilk test. P value is calculated using 1-way ANOVA followed by Tukey post hoc multiple comparisons to compare between the groups (n>3 independent experiments per group); the results are shown as means ± SEM. P values indicated in panels, significant as **p<0.01, ***p<0.001 ****p<0.0001.

**Figure 2.**
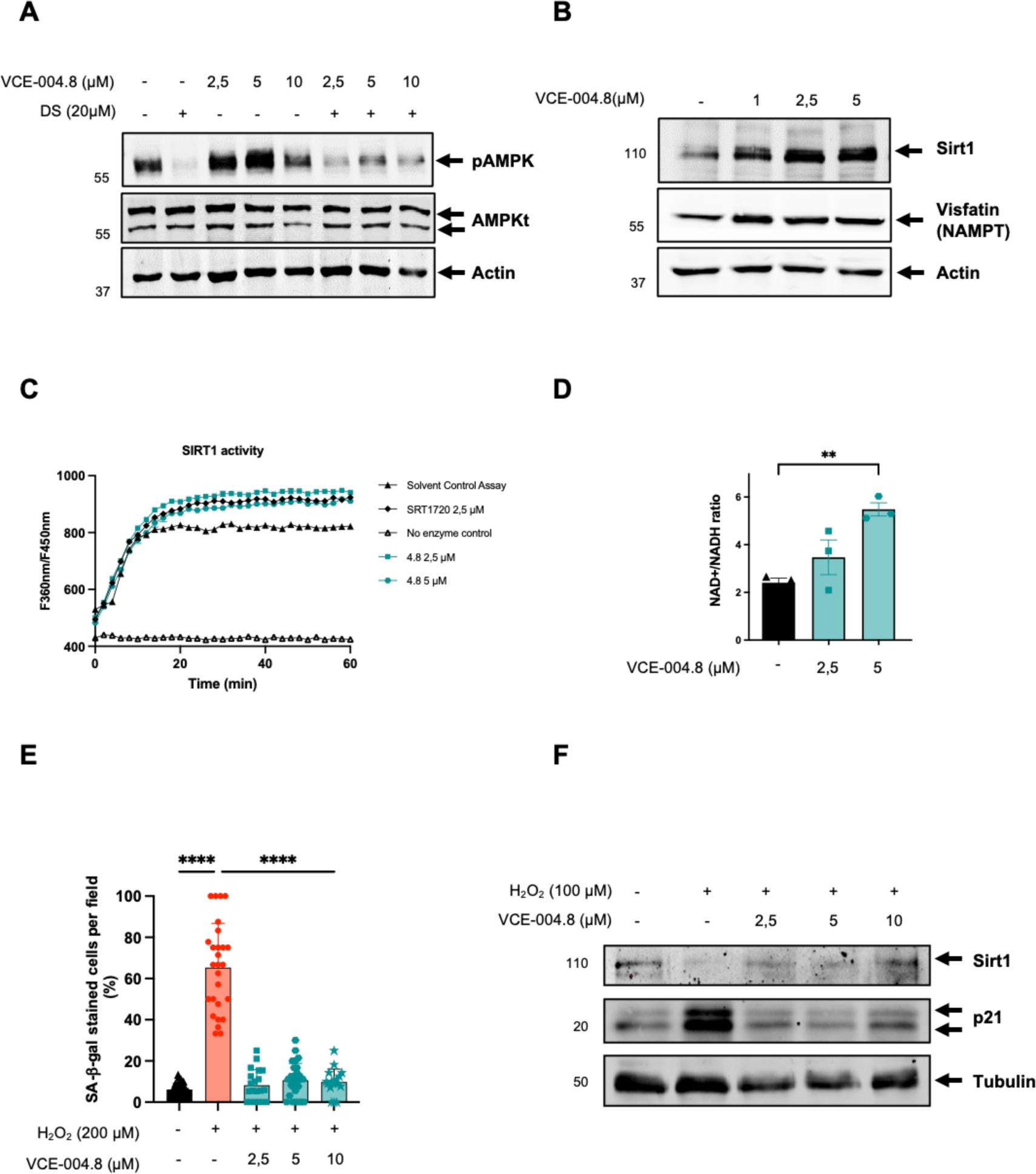
VCE-004.8 activates AMPK/Sirt1 pathway and prevents cellular senescence. **A,** HEK-293T cells preincubated with or without DS for 30 min and stimulated for 3 h with increasing concentrations of VCE-004.8 and the steady state levels of pAMPK total AMPK and actin detected by western blot. **B,** EA.hy296 cells were stimulated with VCE-004.8 for 3h and the expression of Sirt1 and Visfatin (NAMPT) was analyzed by western blot. **C,** VCE-004.8 induces Sirt1 activity measured by a fluorometric assay. SRT1720 was used as a positive control. Data represent the mean ± SD (n = 3). **D,** NAD^+^/NADH ratio was determined in EA.hy296 cells with a NAD^+^ /NADH assay kit. The data passed the normality test performed by the Shaphiro-Wilk test. P value is calculated using 1-way ANOVA followed by Tukey post hoc multiple comparisons to compare between the groups (n=3 independent experiments per group); the results are shown as means ± SEM. P value indicated in panel, significant as **p<0.01. **E,** VCE-004.8 prevents H_2_O_2-_induced senescence measured by SA-β-gal staining in HMEC-1 cells. The data passed the normality test performed by the D’Agostino & Pearson test. P value is calculated using 1-way ANOVA followed by Tukey post hoc multiple comparisons to compare between the groups (n>18 represent areas of cell surface from 3 independent experiments), the results are shown as means ± SD. P values indicated in panel, significant as ****p<0.0001. **F,** HMEC-1 cells were incubated with VCE-004.8 and exposed to H_2_O_2_ and the expression of Sirt1, p21 and tubulin detected by western blot.

### VCE-004.8 activates AMPK/Sirtuin-1 and prevents senescence in endothelial vascular cells

The protective role of the AMP-activated Protein Kinase (AMPK) pathway, a pivotal cellular energy sensor, extends beyond metabolic homeostasis to encompass cardiovascular protection. Vascular AMPK plays a crucial role in regulating endothelial function, redox homeostasis, and inflammation.^20^ Moreover, emerging evidence indicates a functional interplay between AMPK and HIF pathways, contributing to cellular survival adaptation^21^. Thus, we interrogated the participation of AMPK on the pathways activated by VCE-004.8 in endothelial cells. VCE-004.8 induced AMPK phosphorylation and upregulated the expression of Sirt1 and nicotinamide phosphoribosyl transferase (NAMPT), also known as Visfatin, in EA.hy926 cells (Fig. 2A and 2B). A connection between AMPK and Sirt1 has been proposed since AMPK activates Sirt1 through an increase in cellular NAD^+^ level ^22^, while Sirt1 deacetylates the AMPK kinase LKB1, leading to phosphorylation and activation of AMPK^23^. We found that VCE-004.8 increases the enzymatic activity of Sirt1 and the NAD^+^/NADPH ratio in EA-hy926 cells (Fig. 2C and D). Moreover, silencing Sirt1 expression with siRNA prevented the induction of HIF-1α expression by VCE-004.8 (Fig. S3A). To further explore the effect of VCE-004.8 on cellular senescence EA.hy926 cells were treated with H_2_O_2_ and the expression of both SA-β-gal^+^ cells, Sirt1 and p21 proteins analyzed. VCE-004.8 significantly inhibited H_2_O_2_-induced senescence and p21 expression and restored the expression of Sirt1 that was downregulated with the H_2_O_2_ treatment (Fig. 2E and 2F). In addition, pretreatment with dorsomorphin (DS), an AMPK inhibitor, or EX527, a Sirt1 inhibitor, separately or in combination, also inhibited both VCE-004.8-induced EPO-Luc, serving as a surrogated marker of HIF activation (Fig. S3B), and H_2_O_2_-induced cytotoxicity (Fig. S3C). We also investigated eNOS phosphorylation at ser1177, another target of AMPK, and observed that VCE-004.8 induced eNOS phosphorylation (Fig. 3A) as well as NO production (Fig. 3B) in EA.hy926 cells. Interestingly, eNOS phosphorylation was significantly diminished in B55α siRNA cells, providing further evidence for the functional interaction between B55α and AMPK in response to VCE-004.8. Taken together, these findings suggest that in addition to PHD2/HIF-1α activation, the AMPK/Sirt1/eNOS signaling pathways play a significant role in the mechanism of action of VCE-004.8.

**Figure 3.**
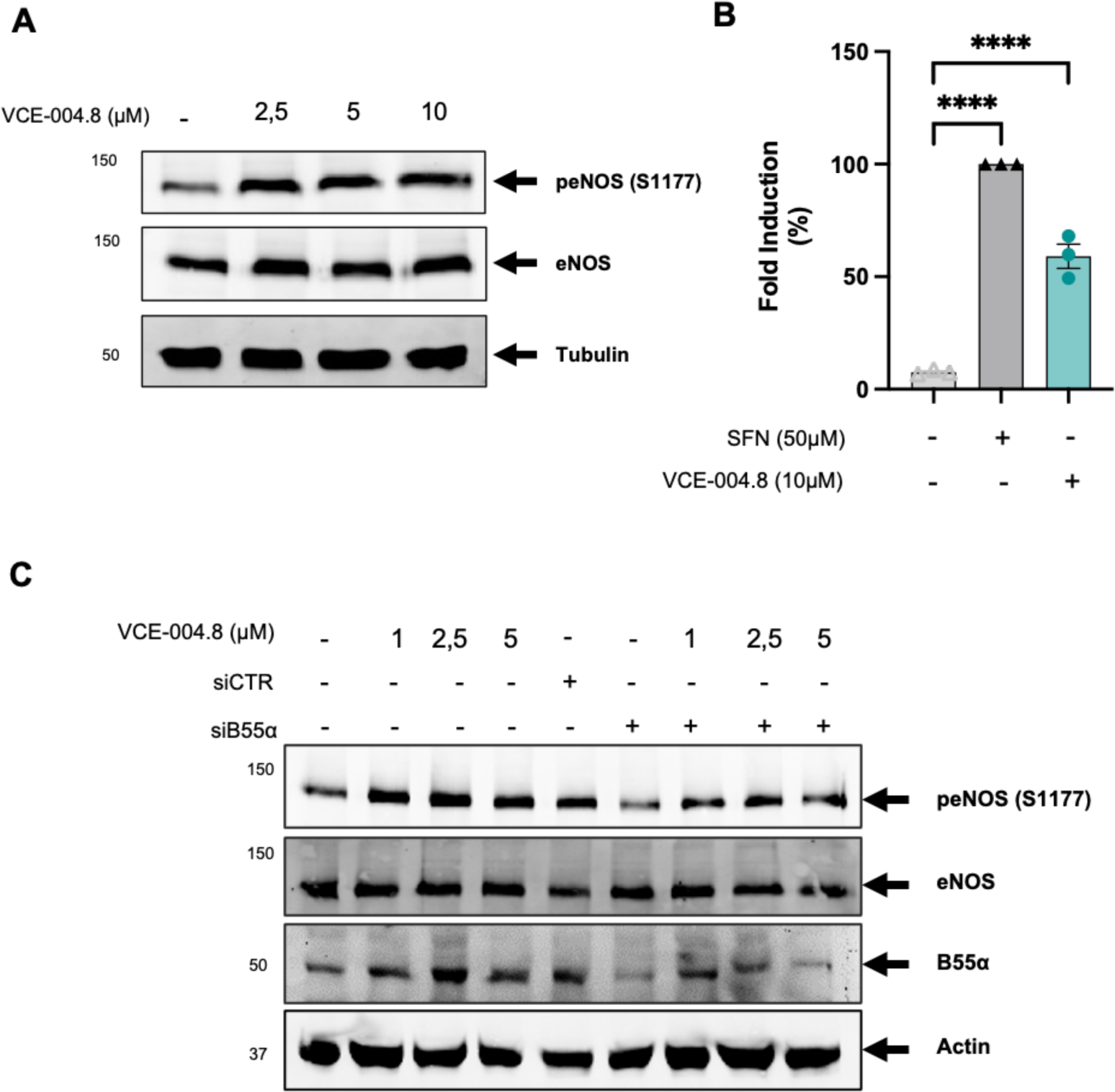
VCE-004.8 induces eNOS activation and NO synthesis in a B55α dependent manner. **A**. EA.hy926 cells were treated with VCE-004.8 for 3 h Representative western blot of three independent experiments is shown. **B,** VCE-004.8 induces NO production. EA.hy296 cells were seeded, preincubated with DAF-2DA and treated with sulforaphane (50 μM) or VCE-004.8 (10μM). The figure indicated the percentage NO induction. The data passed the normality test performed by the Shaphiro-Wilk test. P value is calculated using 1-way ANOVA followed by Tukey post hoc multiple comparisons to compare between the groups (n=3 independent experiments per group); the results are shown as means ± SD. P value indicated in panel, significant as ****p<0.0001. **C,** VCE-004.8 induces eNOS phosphorylation in vascular endothelial cells in a B55α dependent manner. EA.hy296 cells were transfected with siB55α or scrambled siRNAs (siCTR) and 48 h later treated with VCE-004.8 for 3 h. Representative western blot of three independent experiments is shown.

### Angiogenic and arteriogenic effects of VCE-004.8

Vascular regeneration, angiogenesis, arteriogenesis and vasculogenesis are the main objectives for potential vasculogenic therapies for PAD^24^. To evaluate the effect of VCE-004.8 in vascularization and sprouting, we performed an *ex vivo* aortic ring assay. We found that VCE-004.8, as well as VEGF and dimethyloxallyl glycine (DMOG), an enzymatic Pan-PHD inhibitor, induced vascularization *in vitro*, indicated by the formation of new sprouts as well as sprout area (Fig. S4A). Next, to investigate the angiogenic potential of VCE-004.8 in vivo, we conducted a Matrigel plug *in situ* assay. After ten days following the subcutaneous injection of Matrigel into the flank, no signs of angiogenesis were observed in the negative control group (without treatment), in contrast to Matrigel embedded with VEGF (positive control). Oral treatment with VCE-004.8 (10 and 20 mg/kg) clearly increased the formation of functional vessel identified by CD31^+^/αSMA^+^ staining and the proliferative activity of vascular endothelial cells were identified by expression of CD31^+^/Ki67^+^ cells. Moreover, we also confirmed that B55α is also expressed in the new formed vessels (Fig. S4B).

To further explore the effect of VCE-004.8 in a model of critical limb ischemia (CLI) a doble ligation in the femoral arteria was performed in mice and 24 h later animals were treated daily with oral VCE-004.8. After 10 or 28 days of treatment the effect of VCE-004.8 on collateral vessels was investigated by vascular casting and microcomputed tomography (μ-CT) imaging. It is shown in Figure 4A that treatment with VCE-004.8 clearly induced collateral artery formation in the ligated limb compared to the non-ligated control limb, which was not affected by the treatment. These results are in good correlation with the mRNA expression of *Hif1α* and the HIF-regulated proangiogenic genes *Vegfa*, *Epo* and *Hgf* in the gastrocnemius muscle in the affected limb but not in control limb (Fig. 4B) suggesting that the effect of VCE-004.8 is specific to damaged tissues.

**Figure 4.**
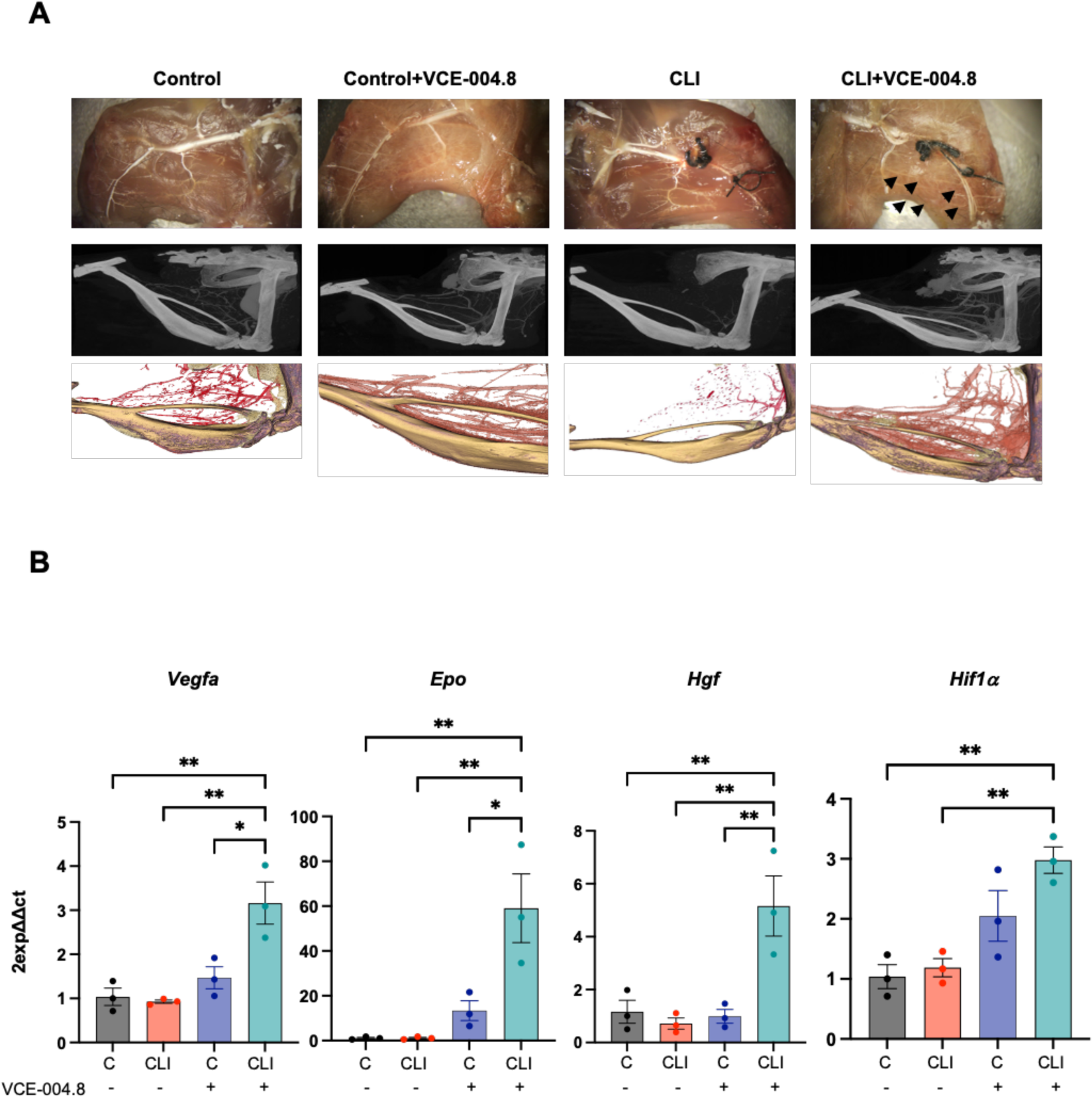
Effects of oral VCE-004.8 in a Critical Limb Ischemia mouse model. **A,** Representative whole-mount images of Microfil vascular cast limbs 10 days after ischemia, Micro-CT reconstruction and their segmentation of arterial vasculature 28 days after femoral artery ligation. Collateral artery growth is indicated by black arrows. **B,** Expression of *Vegf-a*, *Epo*, *Hgf*, and *Hif1α* mRNA in the gastrocnemius after 10 days of ischemia. The data passed the normality test performed by the Shaphiro-Wilk test. P value is calculated using 1-way ANOVA followed by Tukey post hoc multiple comparisons to compare between the groups (n=3 animals per group); the results are shown as means ± SEM. P value indicated in panel, significant as *p < 0.05; **p < 0.01.

### Angiogenic and antifibrotic effects of VCE-004.8 in CLI mice

Reperfusion of ischemic tissues depends on angiogenesis that is required for a proper tissue blood flow^25,26^. Thus, we explored the effect of VCE-004.8 on angiogenesis and vascular density in the gastrocnemius tissue. VCE-004.8 induced angiogenesis measured by the number of vessels (CD31^+^/αSMA^+^) and lumen perimeter vessel per area in the ligated limb without affecting the contralateral limb (Fig. 5A). We also investigated endothelial cells proliferation, a key event in angiogenesis, and found that VCE-004.8 significantly induced cell proliferation as measured by CD31/Ki67 co-staining (Fig. 5B). CD34 is one of the surface markers for endothelial progenitor cells, which plays a major role in angiogenic and vasculogenic activities during neovascularization^27^. Indeed, CD34^+^ cells isolated from peripheral blood have been widely used in experimental therapies including cardiovascular diseases and critical leg ischemia^28, 29^. VCE-004.8 treatment significantly increased the average the of number of CD34^+^/CD31^+^ as well as CD34^+^/CD31^−^ cells in the ligated limb compared with the contralateral limb (Fig. 6A).

**Figure 5.**
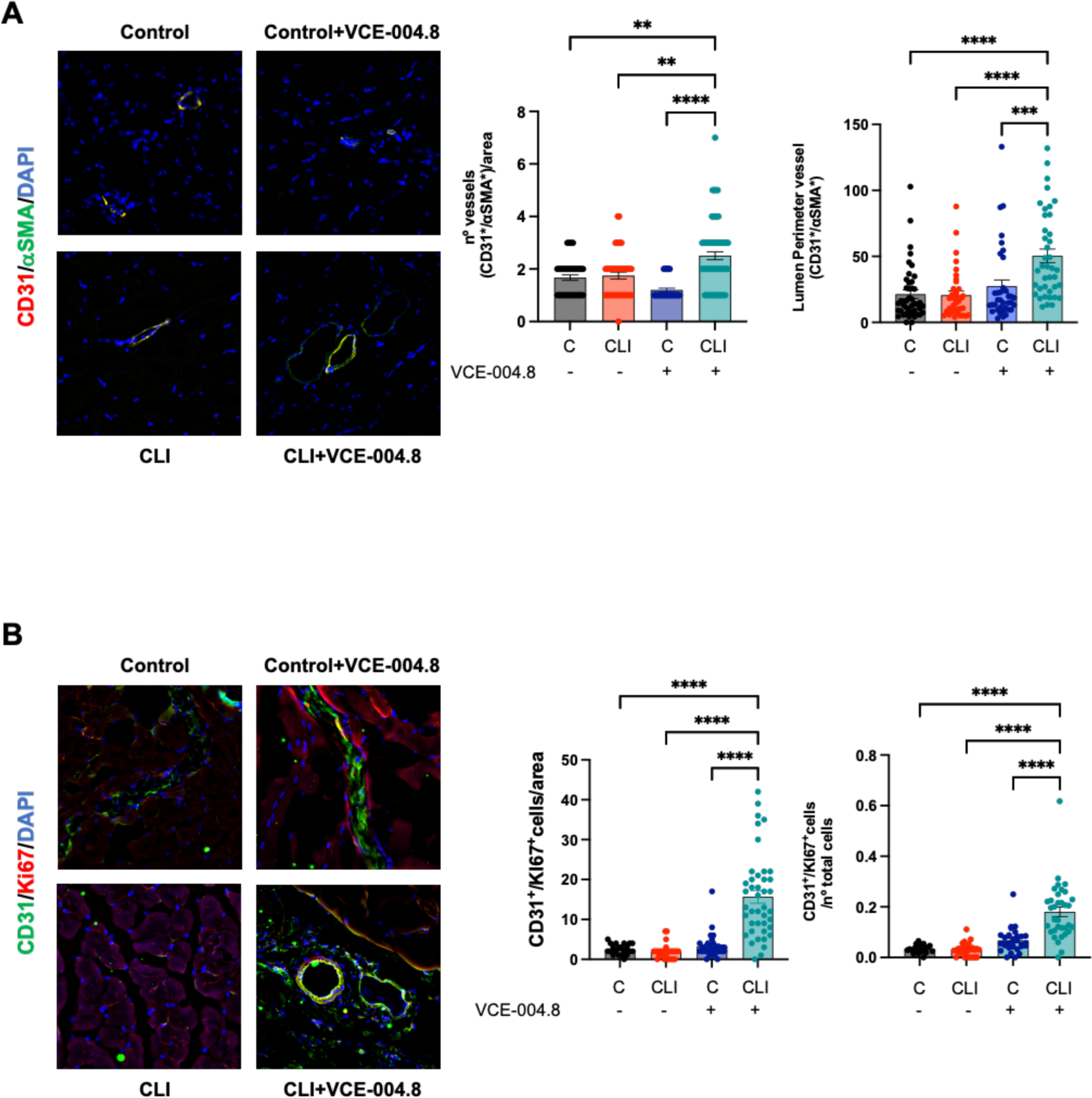
Oral VCE-004.8 enhances ischemia-induced vessel formation and vascular endothelial cells proliferation *in vivo*. **A.** Double immunofluorescence confocal staining of α-SMA (green)/CD31 (red) in gastrocnemius muscles 28 day after ischemia. The immunofluorescence was used to quantify the number of vessel peer area and the perimeter of the lumen vessel in mouse muscle. As the data cannot passed the normality test. P values are calculated using a nonparametric Kruskal-Wallis test followed by Dunn multiple. The results are shown as mean ± SEM. P values indicated in panels, significant as **p < 0.01, ***p<0.001, ****p<0.0001. **B,** The number of CD31^+^/Ki67^+^ positive cells per area and number of CD31^+^/Ki67^+^ cells per total cells were quantified using double immunofluorescence confocal staining of CD31 (green) and Ki67 (red) in gastrocnemius muscles 28 day after ischemia. Significance was determined by one-way ANOVA followed by Tukeýs test. The results are shown as means ± SEM (n=3). P values indicated in panels, significant as ****p < 0.0001.

**Figure 6.**
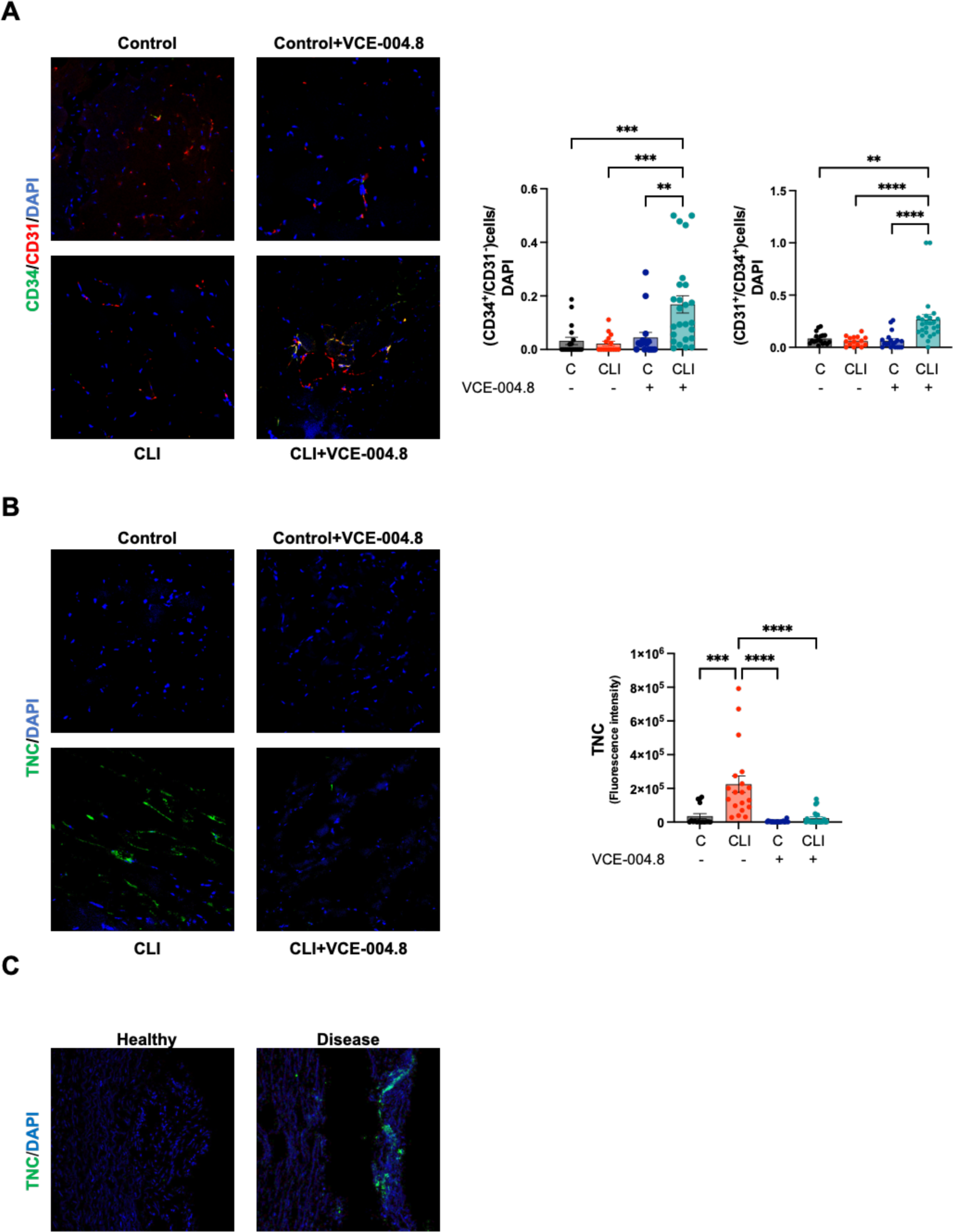
Oral VCE-004.8 enhanced ischemia-induced *de novo* vessel and alleviated fibrosis *in vivo*. **A,** Double immunofluorescence confocal staining of CD34 (green)/CD31 (red) in gastrocnemius muscles 28 days was used to calculate the number of CD34^+^/CD31^−^ cells (immature endothelial vascular cells) and CD34^+^/CD31^+^ (mature endothelial vascular cells). Values are mean ± SEM (n = 3-4). In both quantifications, the significance was determined by one-way ANOVA non-parametric followed by a Kruskal–Wallis test. P values indicated in panels, significant as **p < 0,01; ***p < 0.001; ****p < 0.0001. **B,** Immunofluorescence staining of TNC (green) in gastrocnemius muscles 28 days after ischemia and quantifications. Values are mean ± SEM (n = 3-4), and significance was determined by one-way ANOVA non-parametric followed by Kruskal–Wallis test. P values indicated in panels, significant as ***p < 0.001; ****p < 0.0001. **C,** Representative immunofluorescence staining of TNC in cross-section of healthy and disease human arteries.

Evidence supports the involvement of fibrosis in the pathophysiology of PAD^30^. Fibrosis is associated with various pathological processes in vascular tissues and may contribute to the progression of PAD^31^. Herein, we investigated the expression of Tenascin C (TNC), a marker for fibrosis in both CLI mice and in ex-vivo human samples from PAD patients ^32^. TNC expression was detected in the ligated limb of CLI mice (Fig. 6B) and human aorta from PAD patients (Fig. 6C) and treatment with VCE-004.8 significantly alleviated TNC deposition in the ischemic tissue relative to the contralateral leg (Fig. 6B).

### B55α and Sirtuin 1 expression in CLI mice and human arteries

As demonstrated above, our data revealed a mechanistic link between B55α and Sirt1 expression and activity in vitro. Consequently, we investigated the expression of both proteins in tissues. B55α and Sirt1 expression were found to be downregulated in the ligated limb from CLI mice compared to the contralateral limb (Fig. 7A and 7B), and similar results were observed in the aorta from PAD patients (diseased) compared to healthy tissues (Fig. 7D). VCE-004.8 treatment completely restored the expression of both proteins in the ligated limb of CLI mice. The mRNA expression levels of B55α and Sirt1 confirmed the findings obtained from confocal analysis. Additionally, VCE-004.8 exhibited a tendency to increase the mRNA expression of Sirt1 in the non-ligated limb of CLI mice (Fig. 7C).

**Figure 7.**
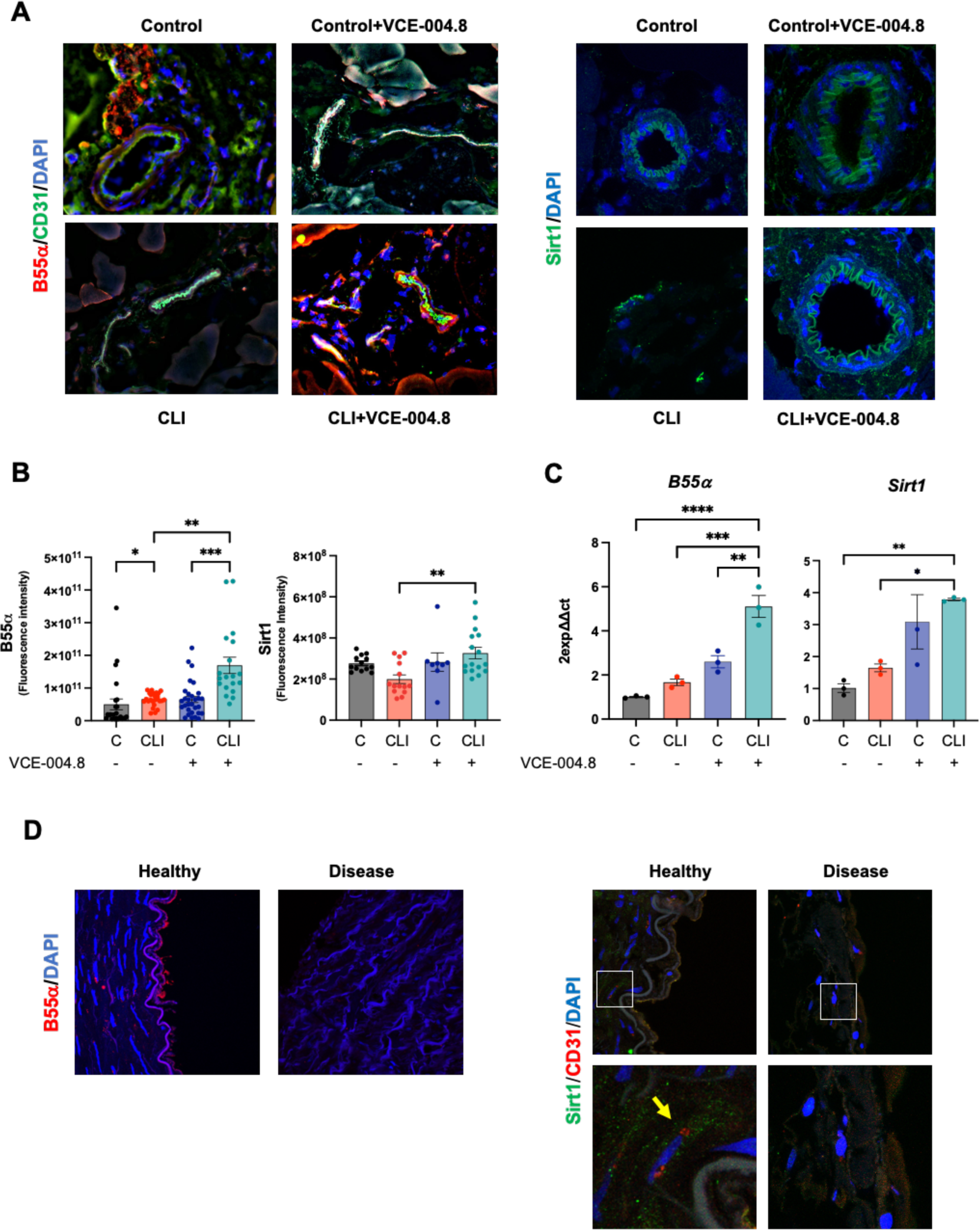
Effects of VCE-004.8 on B55α and Sirt1 expression *in vivo*. **A,** Representative double immunofluorescence confocal staining of B55α (red)/CD31 (green) (left) and immunofluorescence staining of Sirt1(green) (right) in gastrocnemius muscles 10 days after ischemia. **B,** Quantifications of B55α (left) and Sirt1 (right) staining in gastrocnemius muscles 10 days after ischemia. For B55α and Sirt1 quantification, the data do not pass the normality test performed and the significance was determined non-parametric followed by a Kruskal–Wallis test. Values are mean ± SEM (n = 3-4). P values indicated in panels, significant as *p < 0.05 as **p < 0.01; ***p < 0.001. **C,** Expression of *B55α* and *Sirt1* mRNA in mice gastrocnemius at 10 days after femoral ligation. The data passed the normality test performed by the Shaphiro-Wilk test. P value is calculated using 1-way ANOVA followed by Tukey post hoc multiple comparisons to compare between the groups (n=3 animals per group). The results are shown as mean ± SEM. P value indicated in panel, significant as *p < 0.05; **p < 0.01; ***p < 0.001; ****p < 0.0001. **D,** Representative immunofluorescence image of B55α (red) in cross-section of healthy and disease human arteries (left) and representative double immunofluorescence staining of Sirt1 (green) and CD31(red) in cross-section of healthy and disease human arteries (right). Magnified region (white rectangle) showing details of Sirt1 expression (yellow arrow).

### Caveolin 1 is downregulated in CLI mice and human arteries

To investigate the effects of VCE-004.8 on the expression of other markers, we labeled the tissues with αSMA to identify mature vessels. These vessels were subsequently isolated and studied using Laser Capture Microdissection coupled to Mass Spectrometry (Fig. 8A). From the total bulk of proteins (210), eleven proteins involved in the angiogenesis process were identified according to Gene Ontology Annotations from Mice Genomic Identification ^33^. Among them, caveolin 1 (CAV1) is of special interest in our study since its gene is regulated by HIF-1^34^ and it is required for collateral development in CLI mice ^35^ (Fig. 8B). The expression of CAV1 was detected in both the control limb of CLI mice and in healthy human arteries (Fig. 8C and 8E), while it was downregulated in the ligated limb of CLI mice and in arteries affected by PAD (Fig. 8C and 8E). Treatment with VCE-004.8 normalized the expression of CAV1 protein and upregulated its mRNA expression (Fig. 8C and 8D).

**Figure 8.**
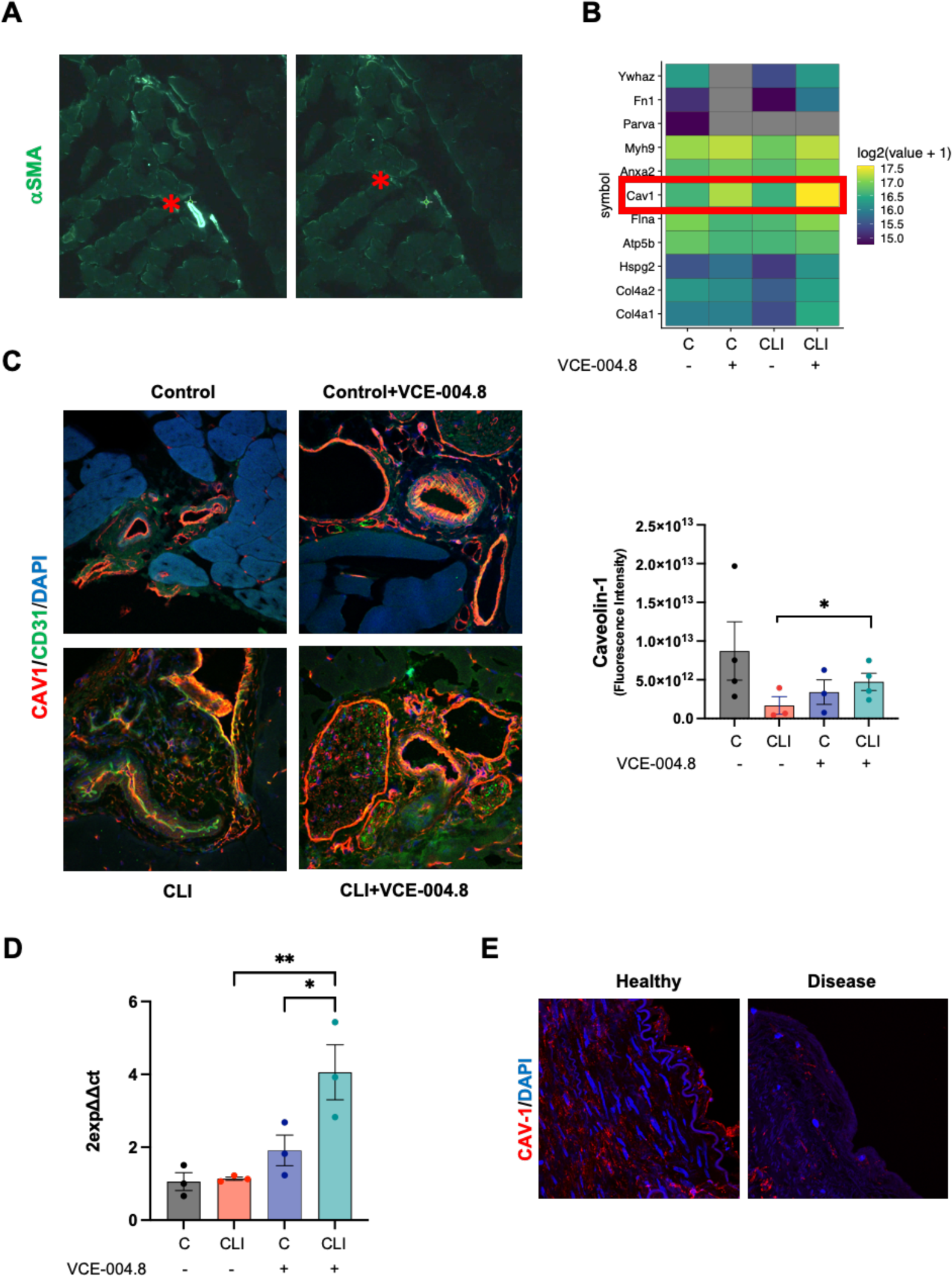
Oral VCE-004.8 induces CAV1 *in vivo*. **A**, Overview of laser capture microdissection. Representative images vessel marked with αSMA in gastrocnemius before (left) and after (right) laser dissection indicated by red asterisk. **B**, Heatmap showing the expression levels for the resulting list of 11 proteins. Color indicates the mean of scaled regularized log transformed expression values. Red square indicates CAV1. **C.** Representatives images of double immunostaining of CAV1(red) and CD31 (green) in gastrocnemius 10 days after ischemia (left) and the CAV1 staining quantification (right). The significance was determined non-parametric followed by a Kruskal–Wallis test. In both quantification, values are mean ± SEM, (n = 3). P values indicated in panels, significant as *p < 0.05. **D,** The expression of *Cav1* mRNA in mice gastrocnemius at 10 days. The data passed the normality test performed by the Shaphiro-Wilk test. P value is calculated using 1-way ANOVA followed by Tukey post hoc multiple comparisons to compare between the groups test (n=3 animals per group). The results are shown as mean ± SEM. Values indicated in panels, significant as *p < 0.05; **p < 0.01. **E,** Representative immunofluorescence staining of CAV1 in cross-section of healthy and disease human arteries.

## DISCUSSION

Effective induction and promotion of angiogenesis have been so far very challenging. A better knowledge of the basic pro-angiogenic players will be crucial to achieve a clinical benefit for PAD patients and other cardiovascular diseases. Therapeutic angiogenesis or vascular regeneration remains an attractive treatment modality for PAD and chronic tissue ischemia. Despite decades of research into angiogenic and cell-based therapies for PAD, their translation to clinical use has been limited. Therefore, there is an urgent need for novel therapeutic angiogenic approaches.

Vascular regeneration strategies include restoration of vascular function and structure that can be partially achieved through HIF activation. While severe hypoxia can be detrimental, leading to cell death and organ failure, mild hypoxia, also known as hypoxia preconditioning, has shown promising therapeutic potential, particularly in the treatment of cardiovascular diseases^36^. Mild hypoxia can be achieved by hypoxia mimetic enzymatic inhibitors of PHDs such are JNJ-42041935, FG-4497, FG-4592 (Roxadustat), TRC160334 (Desidustat), and AKB-4924 ^37^ or by novel PPA2/B55α activators such are Etrinabdione (VCE-004.8) and VCE-005.1^13,38^. Indeed, it has been observed that the enzymatic inhibitor of PHDs, FG-4497, confers protection against the development of atherosclerosis, a pivotal factor in PAD pathogenesis ^39^. Herein, we show the sprouting and angiogenic activity of VCE-004.8 *in vitro* and *in vivo* and its efficacy in CLI mice induced by femoral ligation. Etrinabdione induced endothelial cells proliferation, angiogenic gene expression and prevented fibrosis. One striking and interesting observation is that the effects of Etrinabdione occur only in the affected limbs compared to non-ischemic limbs. A similar observation was found with sodium nitrite therapy and metformin in CLI mice ^40, 41^ and these results agree with our previous finding showing that VCE-004.8 induced angiogenesis and restored damaged brain blood barrier in the affected brain hemisphere of mice with traumatic brain injury but in the contralateral hemisphere ^13^. Thus, we could speculate that the therapeutic benefit of Etrinabdione will be akin to converting severe hypoxia into therapeutic mild hypoxia in the affected limb.

The pathophysiology of atherothrombosis and PAD is complex, and involves many cells, proteins and pathways, giving the possibility to explore multitarget therapeutic strategies, which by covering different angles of the disease may be beneficial for the outcome of patients. Herein we show that Etrinabdione, a multitarget drug candidate in clinical phases, induces HIF-1α expression in endothelial cells through a novel pathway that potentially involves two axes: B55α/PHD2 and B55α/AMPK/Sirt1 signaling that may converge on HIF stabilization. In addition, VCE-004.8 induces the phosphorylation of eNOS at ser1177, which is a target for AMPK. How B55α regulates AMPK needs to be investigated at the molecular level but previous studies have revealed that AMPK signaling regulates the nuclear accumulation of HIF-1α through a complex process involving HDAC5 phosphorylation. This process leads to the export of HDAC5 from the nuclei to the cytosol, where it dissociates the complex formed by the HSP70 chaperone and nascent HIF-1α ^42^. Subsequently, HIF-1α associates with HSP90, facilitating its maturation and relocation into the nucleus. Notably, treatment with CCT018159, an HSP90 inhibitor, partially inhibits VCE-004.8-induced HIF-1α stabilization (Fig. S5). Moreover, Sirt1 interacts with AMPK and HIF-1α, and pharmacological inhibition of Sirt1 impairs HIF-1α nuclear accumulation and the expression of HIF-1α-dependent genes ^43^. Therefore, it is probable that VCE-004.8 requires AMPK to enhance the synthesis of nascent HIF-1α, and B55α/PHD2 to stabilize and translocate this nuclear factor into nuclei. This is supported by our results showing that metformin, an AMPK activator that phosphorylates eNOS ^39^ but did not activate HIF-1α in vascular endothelial cells^44^. However, it is noteworthy that metformin has been shown to promote revascularization in CLI mice independently of HIF-1α ^41^.

In addition, VCE-004.8 is a dual agonist of PPARγ and CB_2_R^14^. PPARγ and CB_2_R are preclinically validated therapeutic targets for vascular inflammation and atherosclerosis ^45,46,47^. For instance, foam cells play a vital role in the initiation and development of atherosclerosis since they are the major sources of necrotic core in atherosclerotic plaques. Interestingly, it has been shown that PPARγ agonists inhibit the formation of macrophage foam cells *in vivo*^48^. Moreover, JHW-015, a CB_2_R agonist, significantly prevented oxLDL accumulation in foam cells and reduced the production of proinflammatory cytokines and the expression of CD36, a scavenger receptor that plays a critical role in the formation of foam cells. In agreement with finding CB_2_R activation by WIN55,212-2 also reduces an OxLDL-induced inflammatory response in rat macrophages^49^. In line with this we have found that VCE-004.8 inhibits the induction of foam cells in OxLDL-treated macrophages (Fig. S6).

On the other hand, VCE-004.8 has also shown efficacy in a murine model of metabolic syndrome induced by high-fat diet. VCE-004.8 showed anti-obesity and anti-inflammatory activity, prevented liver damage and significantly improved glycolipid metabolism^17^. If this activity had translation in humans, we could expect an added benefit in PAD patients with type II diabetes as co-morbidity. The same hold true for coexisting cerebrovascular or coronary diseases since VCE-004.8 showed efficacy in a preclinical stroke model with an excellent therapeutic window and inhibited angiotensin-II signaling and cardiac inflammation and fibrosis^16,18^.

In many cases, diseases with complex physiopathology are very difficult to be treated by monotargeted therapies, and usually requires concomitant therapies (polypharmacy) that have limitations such as therapy adherence and pharmacological interactions. In contrast, multipronged therapies (polypharmacology) may represent potential pharmacological therapies with better clinical benefits after providing a safe profile in humans and an appropriated target engagement as it is the case of reprofiling therapies with small molecules^50^.

## CONCLUSIONS

In conclusion, our results collectively demonstrate that VCE-004.8 is a first-in-class activator of PP2A/B55α, a phosphatase critical for vascular endothelial cell homeostasis through HIF-1 stabilization and activation. Furthermore, VCE-004.8 activates the AMPK/Sirt1/eNOS pathway in a PP2A/B55α-dependent manner, showing potential in preventing endothelial cell damage and senescence, while also promoting arteriogenesis and angiogenesis in CLI mice. Moreover, the supplementary anti-inflammatory properties of Etrinabdione position it as a potential disease-modifying drug, offering clinical benefits to PAD patients with diverse comorbidities

## ACKNOWLEDGMENTS

We acknowledge the Microscopy Advanced Optical Microscopy and the Proteomic Units of IMIBIC for its support.

