## supplemental information for "Etrinabdione (VCE-004.8), a B55α activator, promotes angiogenesis and arteriogenesis in critical limb ischemia"

### SUPPLEMENTAL MATERIAL

Please see the Major Resources Table in the Supplemental Materials

### MATERIAL AND METHODS

**Cell cultures.** EA. hy926 endothelial cells (ATCC® CRL-2922™), HEK-293T (ATCC® CRL-3216™) and NIH-3T3 (ATCC® CRL-1658™) were be cultured in DMEM (Life Technologies, Carlsbad, CA, USA) supplemented with 10% FBS, 2 mM L-glutamine, and 1% (v/v) penicillin/streptomycin at 37 in a humidified 5% CO2 incubator. Human microvascular endothelial cells (HMEC-1) (ATCC® CRL-3243™) were maintained in MCDB131 medium (Life Technologies, Carlsbad, CA, USA) supplemented with 10 ng/mL epidermal growth factor (EGF), 1 µg/mL hydrocortisone, 10 mM glutamine and 10% FBS. For high glucose (HG), cells were be grown in 20 mM, 40 mM or 270 mM glucose and controls were receive 20 mM or 40 mM mannitol or 5 mM glucose.

**Cell viability.** The influence HG medium (serum free), $H_2O_2$ (200µg/ml) or oxLDL (200 µg/ml) in EA. hy926 were assessed by MTT (3-(4,5-dimethylthiazol-2-yl)-2,5-diphenyltetrazolium bromide) assay. Cells were seeded at a density of 10,000 cells/well in 96-well plates. Cells were preincubated with or without VCE-004.8 and inhibitors: HIF1α inhibitor, (YC-1) (#S7958, Selleckchem, Dallas, TX, USA), AMPK inhibitor (Dorsomorphin) (DS) (#S7306, Selleckchem), SIRT1 inhibitor EX527 (#E7034, Sigma, St Louis, MO, USA), HPS90 inhibitor CCT018159 (#385920, Sigma). Then, cells were treated with HG medium (serum free), $H_2O_2$ or OxLDL and then MTT (N6876, Sigma) were added, and survival were determined.

**Luciferase assays and cells transfections.** The mouse NIH-3T3-EPO-Luc cells were stably transfected with the Epo-Luc plasmid that contains three copies of the hypoxia response element (HRE) sequence from the promoter of the erythropoietin gene in the pGL3 vector. NIH-3T3-EPO-luc cells were seeded in 96-well plates and incubated with inhibitors as indicated for 30-60 min before, then VCE-004.8 compound was added. Luciferase activity was quantified using Dual-Luciferase Assay (#E1483, Promega, Madison, WI, USA) after 6h of stimulation. Scramble control oligonucleotide siRNA non-targeting pool (#D-001810) and ON-TARGET plus SMARTpool against B55α (#L-004824) or Sirt1 (L-003540) were purchased from Dharmacon (Waltham, MA, USA).

**Immunocytochemistry analysis.** EA.hy296 cells (2.5 × $10^3$/well) were seeded onto glass coverslips in 24 well plates, then the cells were pre-stimulated with VCE-004.8 at several concentrations for 1h and then incubated for interleukin 1 beta (IL1β) (R&D Systems, Minneapolis, MN, USA) plus tumor necrosis factor alpha (TNFα) (R&D Systems) (for VCAM1 study) or interleukin 6 (IL-6) (R&D Systems) plus TNFα Cells were then washed with PBS (Sigma) and fixed methanol: acetone (Sigma) (1:1) for 8 minutes at -20ºC. After fixation, cells were washed three times with PBS , permeabilized with 0.5% Triton X-100 in PBS at room temperature (RT) for 5 min and blocked in PBS with 3% BSA (Sigma) for 1h. Cells were incubated overnight at 4°C with the following primary antibodies diluted in PBS with 3% BSA: rabbit monoclonal anti- Vascular cell adhesion molecule 1 (VCAM1) (1:100, #ab134047, Abcam, Cambridge, UK) or rabbit polyclonal anti- Zonula Occludens 1 (ZO-1) (1:100, #40-2300, Invitrogen; Carlsbad, CA, USA), Claudin 1 (CLD1) (1:100, #ab15098, Abcam) and PP2A B Subunit Antibody/B55α (B55α) (1:100, #4953T, Cell Signaling, Danvers, MA, USA). Next, cells were washed three times with PBS and incubated with the appropriate secondary antibody for 1 hour at RT in the dark. Finally, coverslips will be mounted with Vectashield Mounting Medium with 4′,6-diamidino-2-phenylindole (DAPI) (Vector Laboratories, Burlingame, CA, USA) for nuclear staining. Images will be acquired using a spectral confocal laser-scanning microscope LSM710 (Zeiss, Jena, Germany) or fluorescence microscope Leica Thunder Imager 3D assay (LEICA, L'Hospitalet de Llobregat, Spain).

**Western blots.** Western blot was performed as previously described [19]. Briefly, after treatments, the cells were washed with PBS and proteins extracted in 50 μL of lysis buffer (50 mM Tris–HCl (pH 7.5), 150 mM NaCl, 10% glycerol, and 1% NP-40) supplemented with 10 mM NaF, 1 mM $Na_3VO_4$, 10 μg/mL leupeptin, 1 μg/mL pepstatin and aprotinin, and 1 μL/mL PMSF saturated. Thirty to forty micrograms of proteins were boiled at 95°C in Laemmli buffer and electrophoresed in 10% SDS/PAGE gels. Immunodetection of specific proteins was carried out by incubation with primary antibody against HIF-1α (1:500 dilution, # ab179483, Abcam), B55α (1:1000 dilution, #5689S, Cell Signaling), phospho-AMPK (1:1000, #2535S, Cell Signaling), total-AMPK (1:1000, #ab80039, Abcam), Sirtuin 1 (1:1000, #8469, Cell Signaling), Visfatin (1:1000, #ab236874,

Abcam), p21 Waf1/Cip1 (p21) (1:1000, #2947, Cell Signaling), HSP 90 (1:1000, #4874, Cell Signaling), α-tubulin (1:5000, #T9026, Sigma) and β-actin (1:10000 dilution, #ab49900, Abcam), overnight at 4 °C. After washing membranes, secondary antibody was added for 1 hour and detected by immunofluorescence system ChemiDoc MP Imaging System (Bio-rad, Hercules, CA, USA).

**Nitric oxide (NO) detection.** The NO production was measured using fluorescent dye DAF-2DA (Merck, Madrid, Spain). EA.hy296 cells were cultured in 96 well plates until 90% confluence and then serum starved overnight. The cells were pre-incubated with 5 μM DAF-2 DA for 30 min at 37 °C in darkness and rinsed with fresh suspension media to remove excess fluorophore and treated with either sulforaphane (50μM) or VCE-004.8 (10μM). After 6 hours, fluorescence signals were measured using IncuCyte zoom (Essen Biosciences, Ann Arbor, MI, USA).

**Senescence assay.** HMEC-1 cells were treated with VCE-004.8 at different doses for 1h, $H_2O_2$ was added for 4h on days 2 and 5 and then culture up to 7 days. SA-β-gal staining was performed according to manufacturer instructions (#9860, Cell Signaling).

**SIRT1 Activity Assay.** SIRT1 activity was determined with the SIRT1 Activity Assay Kit (Fluorometric) (#ab156065, Abcam) that allows the rapid and sensitive evaluation of SIRT1 inhibitors or activators using purified SIRT1, according to the manufacturer's instructions.

**NAD+/NADH Assay.** EA.hy926 ($2 \times 10^6$/well) were seeded and 24h later, the compound was added at several concentrations. After 4h, NAD+ and NADH were measured with a commercially available NAD+ /NADH assay kit (#ab65348, Abcam) according to the manufacturer's protocol.

**THP-1 cell differentiation and foam cell formation**. THP-1 cells were cultured in 24 well plates ($1.5x10^6$ cells/well) and incubated with 160 nM PMA for 24 h, the undifferentiated macrophages were removed, and foam cell formation was induced by incubation of THP-1 derived macrophages with oxLDL (80 μg/ml) or oxLDL + VCE-004.8 for another 48 h.

**Nile red staining**. After foam cell formation, media was aspirated, and the cells were fixed in 4% paraformaldehyde for 20 minutes. Subsequently, cells were rinse with PBS and then stained with freshly diluted Nile red solution (N3013, Sigma)  for 20 min at RT. Thereafter, Nile red solution was removed, and cells were rinse with PBS, and then DAPI and mounting medium were added with Leica Thunder Imager 3D assay (Leica).

**Mouse Aortic Ring Assay.** Male C57BL/6 mice were obtained from Charles River (Barcelona, Spain) and housed with free access to standard food and water under controlled conditions (temperature 20 ◦C (± 2 ºC), 40–50% relative humidity and 12 h light/dark cycle). The thoracic aorta was sectioned into 1 mm long aortic rings and cultured in Opti-MEM (Thermo Scientific) with 100 U/mL penicillin and 100 µg/mL streptomycin overnight. Aortic rings were encapsulated in growth factor reduced Matrigel (Corning, New York, NY, USA) in 24-well plates. The aortic ring was then cultured in Opti-MEM supplemented with 2.5% FBS, 30 ng/mL recombinant human VEGF-A165 (R&D Systems) as positive control or with VCE-004.8 at different concentrations, 2,5 µM and 5 µM in a humidified 37°C, 5% $CO_2$ incubator for 10 days. The rings are fed with growth medium every 2 days. Images were acquired by using a microscope.

**Angiogenesis *in vivo* mouse model**. C57BL/6 male mice 8-10 weeks (n=3 animals per group) were subcutaneously injected with liquid containing Matrigel Gel (Thermo Fischer Scientific) mixed with heparin (0.1 mg/mL) (Ajinomoto Pharma, Tokyo, Japan). For the positive control recombinant human VEGF-A165 (200 ng/mL) (R&D Systems) and fibroblast growth factor 2 (1 µg/mL) (Inmunotools) were added to the gel mix. The Matrigel (500 µL) was injected subcutaneously into flanks of mice. VCE-004.8 (10 or 20 mg/kg) was oral daily administered until experimental endpoint.

**Tissue preparation.** For angiogenesis Matrigel plugs were explanted on day 7 and the connective and adipose tissues surrounding the plug were removed and then fixed in 10% formaldehyde and paraffin embedded for histological analysis. In CLI mouse model, for optimal tissue protection we were carry out perfusion fixation of whole mice with a heparin solution to prevent clotting. First, the mice were anesthetized and injected with 1000 UI of heparin (Sigma), then were perfused in the left ventricle with a vasodilation solution (100 µM adenosine (Sigma),10 µM sodium nitroprusside (Sigma) and 0,05%

BSA in DPBS (Dulbecco's phosphate buffered saline) with $Ca^{2+}Mg^{2+}$ at RT to avoid vasoconstriction. The limb muscles were collected and progressively frozen in isopentane (2-methylbutane) (Sigma) suspended liquid nitrogen to preserve optimal skeletal muscle morphology. Muscle samples were mounted in OCT compound (ProSciTech, Kirwan QLD, Australia) and cut at -21 ºC into 5 µm-thick sections of muscle fibers oriented in transverse direction. All histological assessments were performed on sections that were examined in a blinded fashion.

**Immunofluorescence.** The slides will be fixed at −20 °C in methanol-acetone (Sigma) for 20 min. The slides were boiled for 10 min in sodium citrate buffer (10 mM, pH 6.0) (Sigma) for antigen retrieval. The sections were washed three times in PBS containing 0.1% Triton X-100 (Sigma). Nonspecific antibody-binding sites were blocked for 1 h at RT with 3% BSA in PBS. Next, the sections were incubated overnight at 4ºC with the following primary antibodies diluted in PBS with 3% BSA: anti-α-Smooth Muscle Actin (α-SMA) (1:500, #53-9760-80; Invitrogen), anti-rabbit or -rat cluster of differentiation 31 (CD31), (1:100, #ab28364, Abcam or 1:100, #14-0311-85, Invitrogen), anti-rabbit Ki67 (1:100, #ab15580, Abcam), anti- rabbit PP2A/B55α (1:100, #4953T, Cell Signaling), rat anti-CD34 (1:100, #553731 BD Biosciences, San Jose, CA, USA) anti-rat Tenascin (TNC) (1:100 dilution,#MAB2138, R&D system), anti-mouse Sirtuin-1 (SIRT1) (1:50, #8469, Cell Signaling), and anti-rabbit Caveolin 1 (CAV1). (1:1000, #3267, Cell Signaling). The next day sections were washed three times during 10 min with a wash buffer and incubated in darkness at RT for 1h with the appropriated secondary antibody. The slides were then mounted using Vectashield Antifade Mounting Medium with DAPI (Vector Laboratories, Burlingame, CA, USA). All images were acquired using a spectral confocal laser-scanning microscope LSM710 (Zeiss) with a 20 x /0.8, 25, 40, or 63 ×/0.8 Plan-Apochromat oil immersion lens and quantified in randomly chosen fields using ImageJ software (http://rsbweb.nih.gov/ij/).

**Quantitative reverse transcriptase PCR analysis.** Total RNA was isolated from gastrocnemius muscle using QIAzol lysis reagent and the RNeasy Lipid mini kit (Qiagen, Hilden, Germany) according to the manufacturer´s instructions. For quantitative reverse transcriptase-PCR assays, total RNA (1 µg) was retrotranscribed using the iScript cDNA Synthesis Kit (Bio-Rad) and the cDNA was analyzed by real-time PCR using the iQTM SYBR Green Supermix (Bio-Rad) and a CFX96 Real-time PCR Detection System (Bio-

Rad). Gene expression of several markers indicated in the table: *VegfA*, *Epo, Hgf*, *Hif1α*, *B55α*, *Sirt1* and *Cav1* were quantified using the 2-ΔΔCt method and the percentage of relative expression against Sham group will be represented. Glyceraldehyde-3-Phosphate Dehydrogenase (*Gapdh*) gene were used to standardize mRNA expression in each sample. The primers used in this study were acquired in Eurofins (Luxembourg) and are listed in the following table:

| Genes | Forward (5”-3”) | Reverse (5”-3”) |
|---|---|---|
| ***VegfA*** | CTGCTGTAACGATGAAGCCCTG | GCTGTAGGAAGCTCATCTCTCC |
| ***Epo*** | GACAAAGCCATCAGTGGTCTACG | GCAGAAAGTATCCACTGTGAGTG |
| ***Hgf*** | GTCCTGAAGGCTCAGACTTGGT | CCAGCCGTAAATACTGCAAGTGG |
| ***Hif1α*** | CCTGCACTGAATCAAGAGGTTGC | CCATCAGAAGGACTTGCTGGCT |
| ***B55α*** | GAGACAAAAGACCAGAAGGCTATA | ATTCTCCGTGGACTGGCTTCCA |
| ***Sirt1*** | GGAGCAGATTAGTAAGCGGCTTG | GTTACTGCCACAGGAACTAGAGG |
| ***Cav1*** | CACACCAAGGAGATTGACCTGG | CCTTCCAGATGCCGTCGAAACT |
| ***Gapdh*** | GGCAAAGTGGAGATTGTTGCC | AAGATGGTGATGGGCTTCCCG |

**Proteomic sample preparation.** Samples were digested with advanced iST(in-Stage-Tip) double digestion using Trypsin and LysC as enzymes. Then, the samples were suspended by the IMSMI staff in a phase A solution and measured using microfluorimetry (Qubit™ Protein Assay). Then, they were analyzed by LC-MS/MS using DDA-PASEF mode on the EvosepOne-TIMSTOF-Flex Clinical Proteomics Platform IMSMI. During acquisition, the samples were identified in real time by PaSER software (https://cutt.ly/L0hMYde) to ensure the correct data acquisition (quality control). All samples were loaded in triplicate. Later, the samples were analyzed using LC run (60SPD) and standard DDA-PASEF method. Fragpipe software and the MSstats shiny application were utilized. In total, 36 runs were performed, including 12 HeLaQC analysis (external QC) to system conditioning and system monitoring, 4 pool QCs, 20 DDA-PASEF analyses containing the triplicates from each sample.

**LC-MS analysis.** Clinical Proteomics Platform of IMSMI was used. It consists of an EvosepOne nanoLC (Evosep, Odense, Denmark, 2021) coupled to a TIMSTOF-Flex (Bruker Daltonics, Bremen, Germany, 2022). For the DDA-PASEF method, each top N acquisition cycle consisted in ten PASEF MS/MS. The collision energy was linearly decreased from 59 eV at 1/K0 = 1.60 Vs cm−2 to 20 eV at 1/K0 = 0.60 Vs cm−2. The accumulation and ramp times were set to 100 ms. Singly charged precursors were excluded from fragmentation using a polygon filter in the (m/z, 1/K0) plane. Additionally, all precursors that reached the target value of 20,000 were excluded for 0.4 min. Precursors were isolated using a Q window of 2 m/z for m/z <700 and 3 m/z for m/z >800.

**Data analysis.** Acquired DDA-PASEF runs of all samples present in the study were used to perform the entire analysis including identification and quantification of each replicate of each sample. To achieve this, we used Fragpipe v19.1 (https://fragpipe.nesvilab.org) with default parameters was used. MSStats shiny package software was used to perform the analysis between user's defined groups. In summary, the entire sample dataset (with exception of samples QC replicates) with all conditions/groups (4 different groups) were used for the comparisons. A log2 data transformation was done and the normalization of transformed data was done by medians equalization. All features were used during the analysis considering a model-based imputation (internally calculated by MSstats) and all NaN values were censored. T-test analyses were performed and a table with potential candidate results was generated. The mass spectrometry proteomics data have been deposited to the ProteomeXchange Consortium via the PRIDE partner repository with the dataset identifier PXD050360.

### SUPPLEMENTAL FIGURES

**Figure S1. VCE-004.8 protects endothelial vascular cell**. EA.hy926 cells were pre-incubated with different concentrations of VCE-004.8 during 1h  and then exposed to either Glucose (270mM) or OxLDL (100 μM). Cell viability was calculated by MTT assay. The data passed the normality test performed by the Kolmogorov-Smirnov test. P value is calculated using 1-way ANOVA followed by Tukey post hoc multiple comparisons to compare between the groups (n>4 independent experiments per group), the results are shown as mean ± SEM.

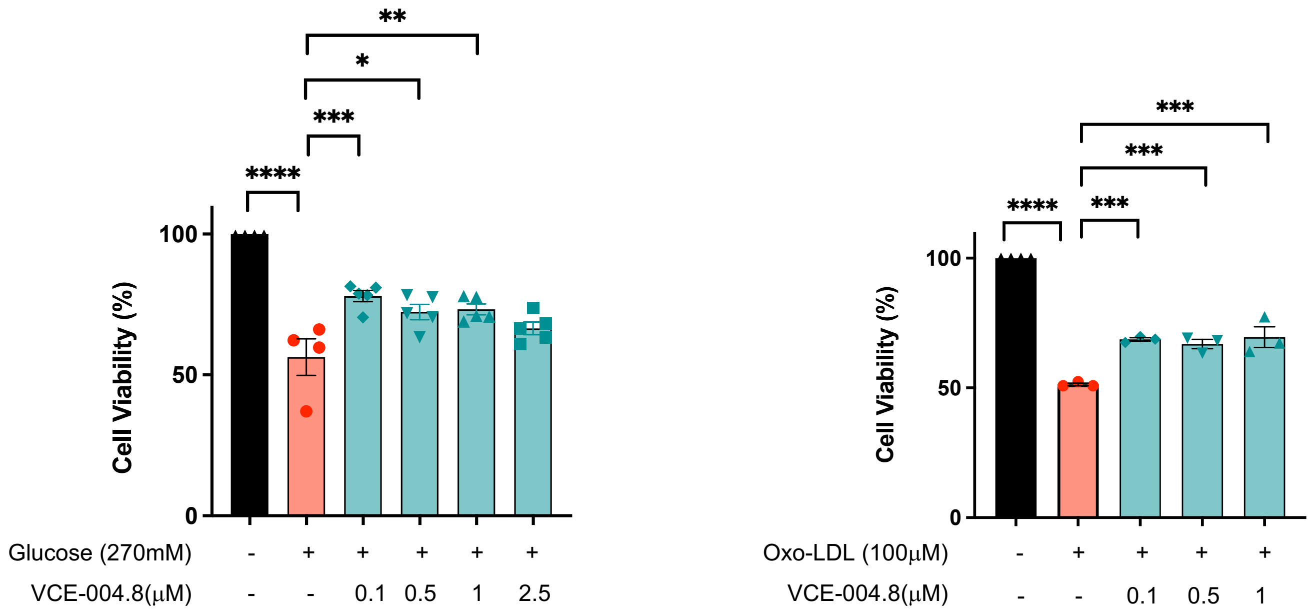

**Figure S2. VCE-004.8 alleviates inflammation of vascular endothelial cells.** Representative images of VCAM, ZO-1, CLD1 and B55α expression in EA.hy926 cells stimulated for 24h with proinflammatory cytokines and their quantifications. The data passed the normality test performed by the Shapiro Wilk test. P value is calculated using 1-way ANOVA followed by Tukey post hoc multiple comparisons to compare between the groups (n=3 independent experiments per group). The results are shown as mean ± SEM. P values indicated in panels, significant as **p<0.01, ***p<0.001 ****p<0.0001.

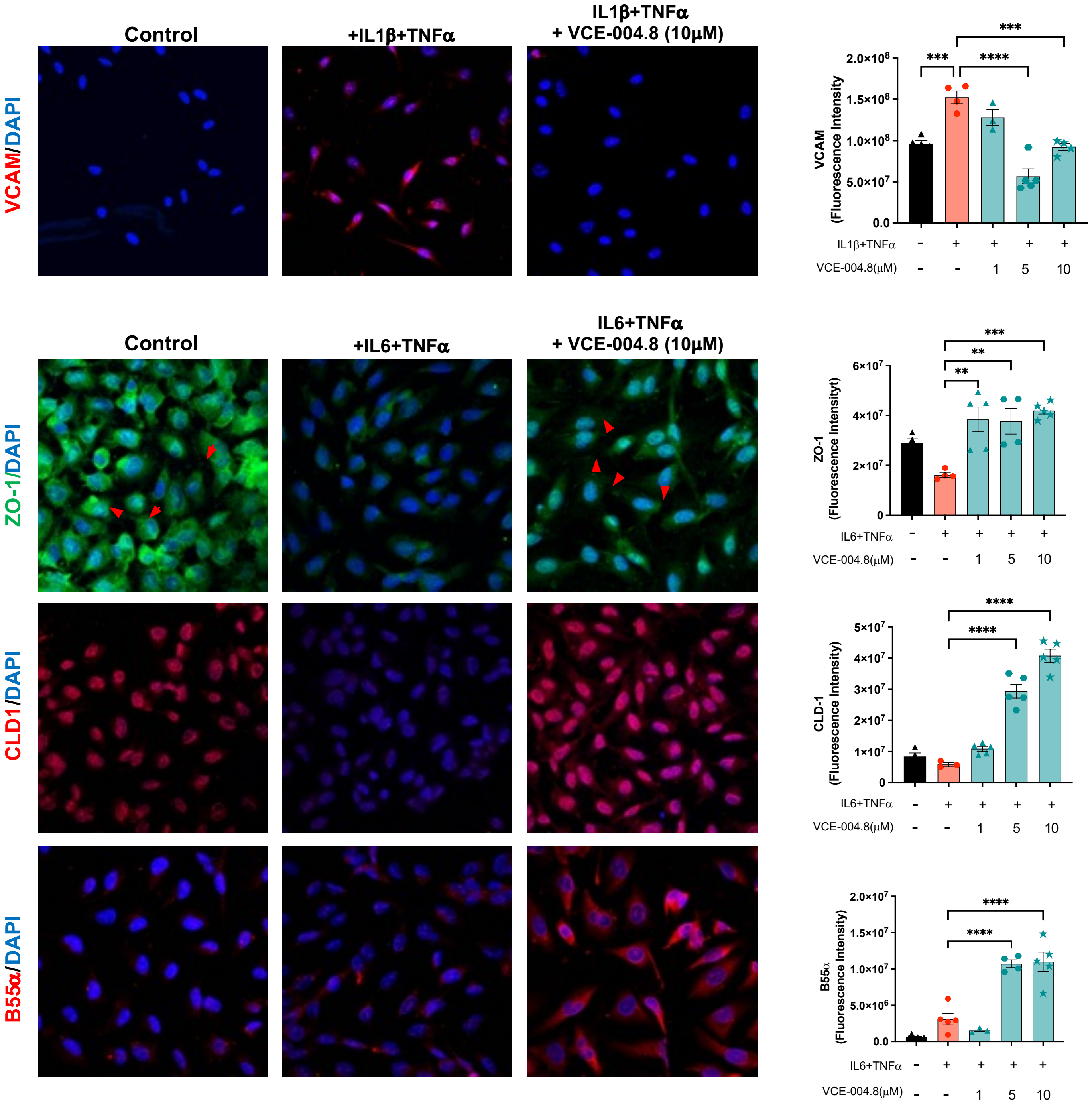

**Figure S3. VCE-004.8 activates HIF-1α and prevents $H_2O_2$-induced cytotoxicity through AMPK/Sirt1**. **A,** HEK-293T cells were transfected with siSIRT1 or scrambled siRNA and treated with VCE-004.8 for 3 h. **B,** NIH-3T3-EPO-luc cells were treated with VCE-004.8 in the absence and the presence of AMPK and Sirt1 inhibitors for 6 h and luciferase activity measured and expressed as percentage of activation considering VCE-004.8 as 100% activation. Data represent the mean ± SD (n =2-7). The significance was determined non-parametric followed by a Kruskal–Wallis test. P values indicated in panels, significant as *p<0.05, **p<0.01. **C,** EA.hy926 cells were preincubated either with DS or Ex527 and treated with VCE-004.8 during 1h before exposure to $H_2O_2$ for 24 h. Cytotoxicity was measured by the MTT method. The data passed the normality test performed by the Shapiro- Wilk test. P value is calculated using 1-way ANOVA followed by Dunnett's post hoc multiple comparisons to compare between the groups. Data represent the mean ± SD (n =3-12). P values indicated in panels, significant as **p<0.01, ****p<0.0001.

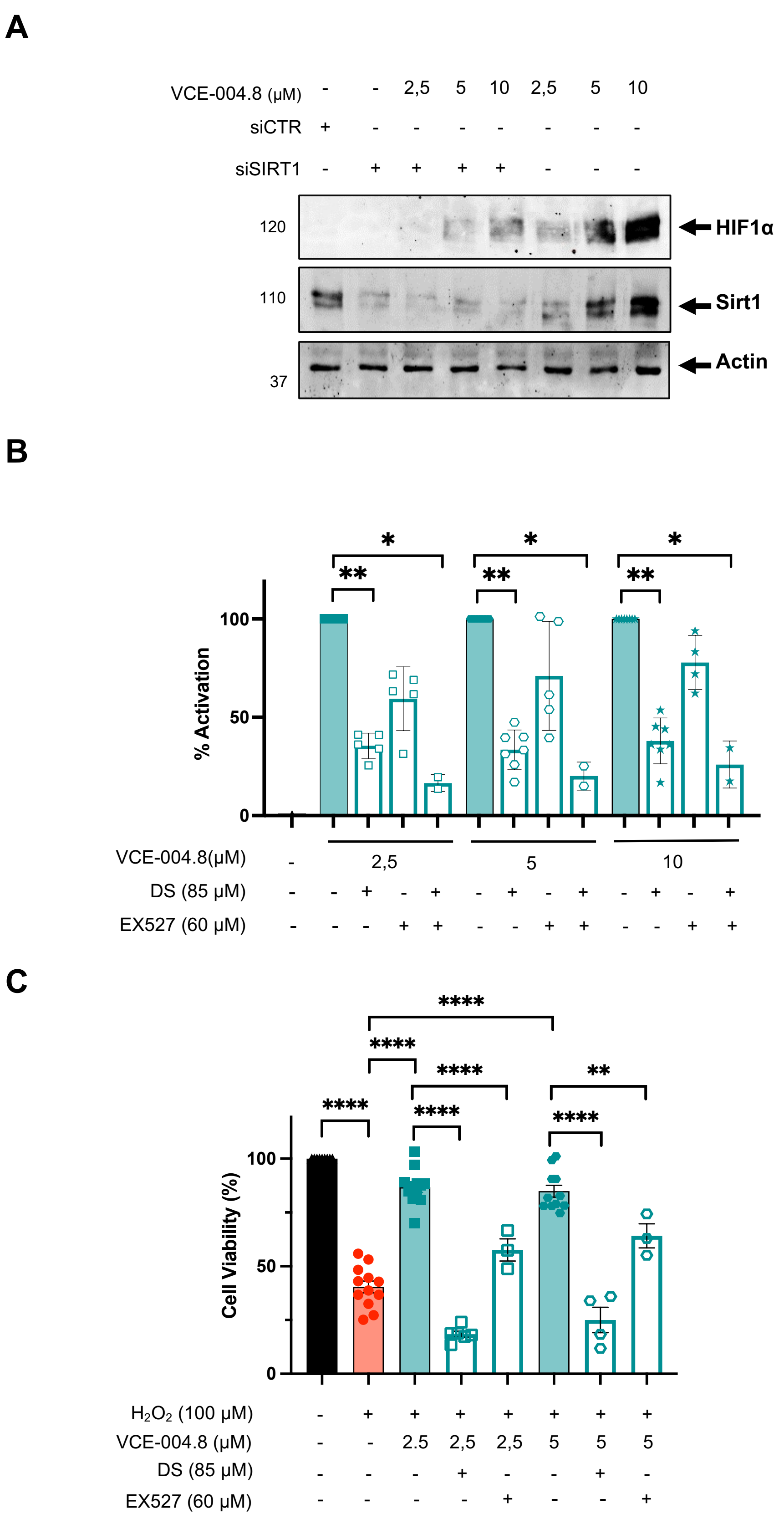

A
VCE-004.8 (μM) - - 2,5 5 10 2,5 5 10
siCTR + - - - - - - -
siSIRT1 - + + + + - - -
120
110
37
HIF1α
Sirt1
Actin
B
% Activation
100
50
0
*
**
VCE-004.8(μM) - 2,5 5 10
DS (85 μM) - - + - + - + - + - + - +
EX527 (60 μM) - - - + + - - + + - - + +
C
Cell Viability (%)
100
50
0
****
**
$H_2O_2$ (100 μM) - + + + + + + +
VCE-004.8 (μM) - - 2.5 2,5 2,5 5 5 5
DS (85 μM) - - - + - - + -
EX527 (60 μM) - - - - + - - +

**Figure S4. VCE-004.8 induces angiogenesis *in vitro* and *in vivo* A,** Representative images of aortic ring assay. Red arrows indicate sprouts and red lines delimit sprouting area. **B,** Macroscopic (top) and confocal images of immunofluorescence staining of Matrigel sections of double immunofluorescence staining of α-SMA (green)/CD31 (red), CD31 (green)/Ki67(red), B55α (green)/CD31(red). For α-SMA staining quantification, the data passed the normality test performed by the Kolmogorov-Smirnov test. P value is calculated using one-way ANOVA followed by Tukey test compare between the groups (n=3). The results are shown as mean ± SEM. For CD31 staining quantification, the significance was determined non-parametric followed by a Kruskal–Wallis test (n=3). The results are shown as mean ± SEM. Double staining CD31/Ki67 was used to quantify vascular endothelial cells proliferation per area ($CD31^+/Ki67^+$ cells). The significance was determined non-parametric followed by a Kruskal–Wallis test (n=3). The results are shown as mean ± SEM. P values indicated in panels, significant as *p<0.05, **p<0.01, ***p<0.001.

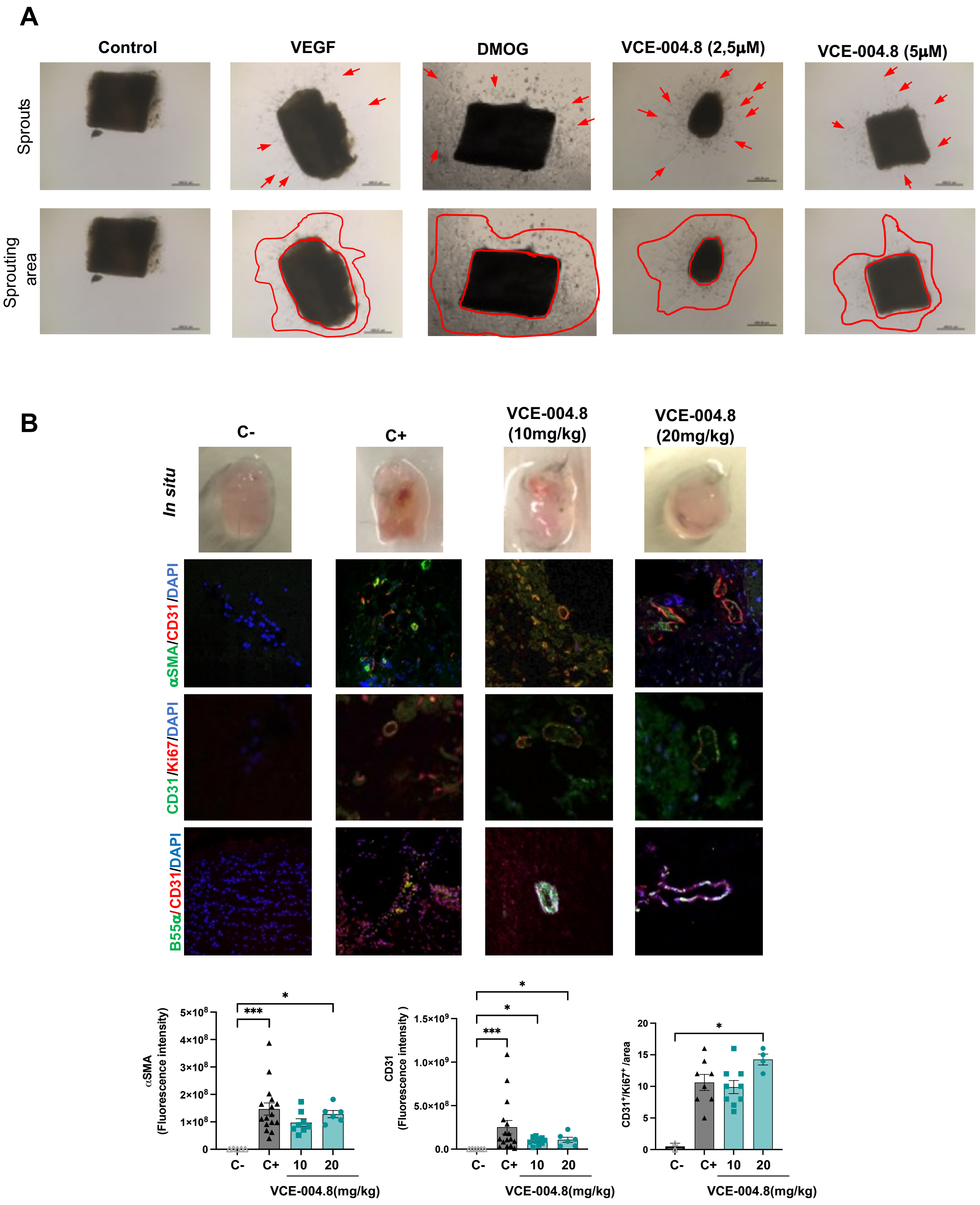
A
Control
VEGF
DMOG
VCE-004.8 (2,5µM)
VCE-004.8 (5µM)
Sprouts
Sprouting area
B
C-
C+
VCE-004.8 (10mg/kg)
VCE-004.8 (20mg/kg)
In situ
αSMA/CD31/DAPI
CD31/Ki67/DAPI
B55α/CD31/DAPI
αSMA (Fluorescence intensity)
CD31 (Fluorescence intensity)
CD31+/Ki67+/area
VCE-004.8(mg/kg)
C-
C+
10
20

**Figure S5**. **The SP90 inhibitor CCT018159 impaired VCE-004.8-induced HIF-1α expression**. EA.hy926 cells were preincubated with **CCT018159** and treated with VCE-004.8 during 3 h. Western blots are representative of 3 independent experiments.

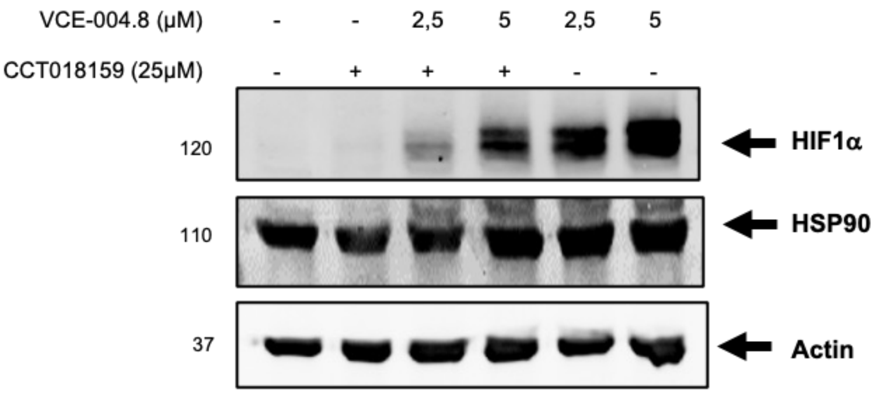

**Figure S6**. **VCE-004.8 prevented oxLDL-induced foam cells measured by Nile Red staining of lipid droplets**. The data passed the normality test performed by the Shapiro-Wilk test. P value is calculated using 1-way ANOVA followed by Dunnett's post hoc multiple comparisons to compare between the groups. Data represent the mean ± SD (n =3-6). P values indicated in panels, significant as ***p<0.001 ****p<0.0001.

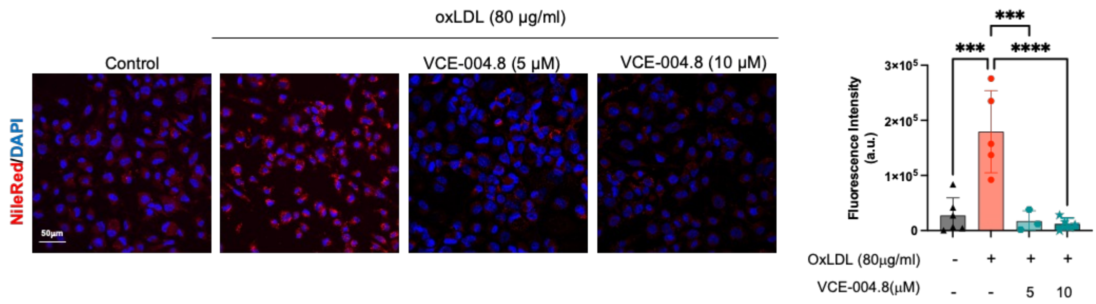
